# Rhythmic tapping to a moving beat: motion kinematics overrules motion naturalness

**DOI:** 10.1101/2023.03.13.532241

**Authors:** Oswaldo Pérez, Sergio Delle Monache, Francesco Lacquaniti, Gianfranco Bosco, Hugo Merchant

## Abstract

Beat induction is the cognitive ability that allow humans to listen to a regular pulse in music and move in synchrony with it. Although auditory rhythmic cues are known to induce more consistent synchronization than flashing visual metronomes, this asymmetry can be canceled out by visual moving metronomes. Here, we investigated whether the naturalness of the visual motion or its kinematics could provide a synchronization advantage over flashing metronomes. Subjects tap in sync with visual isochronous metronomes defined by vertically or horizontally accelerating and decelerating motion, either congruent or not with natural gravity, and then continue tapping with no metronome. We found that motion kinematics was the predominant factor determining rhythm synchronization, as accelerating moving metronomes in either cardinal direction produced more precise and predictive tapping than decelerating or flashing conditions. Notably, a Bayesian observer model revealed that error correction during tapping synchronization and regression towards the mean in accuracy during tapping continuation in the absence of external cues are optimal control strategies independently of the moving properties of the visual metronomes. Our results support the notion that accelerating visual metronomes convey a strong sense of beat as seen in the cueing movements of an orchestra director.

## Introduction

Beat induction is the cognitive ability that allow humans to listen to a regular pulse in music and to move in synchrony with it. Critically, without beat induction there is no music perception and, hence, is considered a universal human trait (Honing, 2012; Honing et al., 2012; Merchant et al., 2015a). A rather useful paradigm to investigate beat induction is the Synchronization-Continuation Task (SCT), in which subjects tap in sync with periodic sensory cues that generate an internal reference interval (synchronization epoch) and keep tapping after the metronome is extinguished using this internal beat representation (continuation epoch) (Wing, 2002; Balasubramaniam et al., 2021). Performance in this task shows two main timing features: a linear increase in variability of produced intervals with the mean that is a form of Weber’s law (called scalar property) (Gibbon et al., 1997; Merchant et al., 2008b; García-Garibay et al., 2016) and an over and underestimation of intervals for shorter and longer intervals respectively (termed regression towards the mean or bias effect) (Woodrow, 1934; McAuley and Jones, 2003; Perez and Merchant, 2018). In addition, subjects use an error correction mechanism that maintains tap synchronization with the metronome, since a longer produced interval tends to be followed by a shorter interval and the other way around, to avoid error accumulation and losing the metronome (Repp, 2005; Iversen et al., 2015). In contrast, during continuation there is a drift in the produced duration (Madison, 2001; Collier and Ogden, 2004). Functional imaging and neurophysiological studies have shown that beat induction and entrainment depends on the cortico-thalamic-striatal involving the medial premotor areas (MPC: SMA and preSMA) and the putamen (Grahn and Rowe, 2009; Merchant et al., 2011, 2013; Bartolo et al., 2014; Sánchez-Moncada et al., 2022). The neural clock flexibly represents the internal beat in the cyclical resetting of cell population states within MPC (Crowe et al., 2014; Merchant et al., 2015b; Merchant and Bartolo, 2018; Gámez et al., 2019; Betancourt et al., 2022).

It is widely known that rhythmic tapping is more accurate and consistent when cued by auditory than flashing visual metronomes (Chen et al., 2002; Repp and Penel, 2002, 2004; Merchant et al., 2008b). The superiority of audition across studies supported the notion that the auditory system is specialized for time, while vision is dedicated for spatial processing (Welch and Warren, 1980; Kubovy, 1988; Guttman et al., 2005). Indeed, recent hypothesis accentuate the role of the audiomotor system in beat perception and entrainment (Honing and Merchant, 2014; Merchant and Honing, 2014; Patel and Iversen, 2014; Lenc et al., 2021). Nevertheless, the auditory–visual asymmetry can be canceled out by using visual moving metronomes (Hove et al., 2010), especially with naturalistic stimuli such as videos of a bouncing ball with changes in speed (Hove et al., 2013; Iversen et al., 2015) or with acceleration profiles that are congruent with the effects of gravity (Gan et al., 2015). The hypothesis behind these observations is that visual motion may engage the time to collision mechanism in parietal cortex used for moving target interception or for collision avoidance (Merchant et al., 2003, 2004a, 2004b; Merchant and Georgopoulos, 2006; Li et al., 2022). Thus, the parieto-premotor system recruited for encoding time-to-contact for single events, could also be involved in the rhythm internalization of periodic collision points, efficiently driving the motor system for beat-based timing (Mendoza and Merchant, 2014; Iversen et al., 2015; Comstock et al., 2018). In addition, the degree of congruency between the visual motion and the motor response can influence rhythm synchronization, as tapping movements aligned with the direction of the visual motion accomplish better synchronization than when the tapping and visual motion direction are incongruent (Hove et al., 2010). These findings are in line with the notion that gravitational cues can facilitate some perceptual functions, like interpreting the causality and the naturalness of object motion or discriminating pendular motion (Pittenger, 1990; Kim and Spelke, 1992; Bingham et al., 1995; Twardy and Bingham, 2002), and can lead to an advantage in manual interceptions, by engaging an internal representation of gravity in the vestibular cortex (Indovina et al., 2005; Lacquaniti et al., 2013; Delle Monache et al., 2021). Interestingly, this internal representation of gravity can be engaged even by the mere object kinematics displayed on a computer screen, especially over a pictorial background providing scaling cues to real world metrics (Moscatelli and Lacquaniti, 2011; La Scaleia et al., 2015; Delle Monache et al., 2017, 2019; Ceccarelli et al., 2018).

No studies have used naturalistic videos to determine the effects of Earth’s gravity on beat induction. Thus, we designed a SCT cued by either vertical or horizontal accelerated objects’ motion, rendered over pictorial background scenes providing elements of naturalness. By pairing upward / downward motion to accelerated / decelerated motion, vertical motion was either compatible or not with Earth’s gravity effects. Thus, downward accelerations and upward decelerations can be considered congruent with gravity effects, whereas downward decelerations and upward accelerations are not. In addition, we also used horizontal motion, where all combinations of motion direction (leftward | rightward) and acceleration (positive | negative) could be considered plausible. In principle, this experimental design can dissect the effects of the target kinematics per se from those of the target motion naturalness on the subjects’ ability to reproduce the temporal intervals cued by the moving stimuli.

Importantly, we developed a Bayesian observer model that fully explained the changes in time precision, accuracy, and correlation in produced intervals in the tapping sequence. By using this framework of statistical inference, we found that subjects used different strategies to optimally switch between the sensory guided tap synchronization and the internally driven continuation, using an error correction and the regression towards the mean mechanism, respectively. Overall, the present findings describe profound effects of the acceleration/deceleration profile of moving metronomes on rhythmic tapping, but strikingly very little effects of natural gravity compared to arbitrary motion conditions. These results go in line with the cardinal movements with sharp acceleration profiles and abrupt changes in direction used by a conductor to define the beat of an orchestra.

## Results

Sixteen subjects performed a modified version of the SCT (Figure 1, see Methods). Briefly, participants produced five intervals by tapping on a touchscreen in sync with cueing stimuli presented alternatively between two positions (synchronization epoch), and then continued tapping with the same tempo for other five intervals in the absence of the visual metronome (continuation epoch) (Figure 1A). Tapping was performed also alternatively between two positions in the bottom right portion of the touchscreen. We used two stimuli and two response locations to generate a SCT that simulate more naturalistic situations, such as a drummer playing the bongos, congas, or timbales (Figure 1B and 1C). In one variation of the SCT, eight different motion conditions were used to define the intervals: two vertical (up and down) and two horizontal (right and left) directions, each with either accelerating (1G) or decelerating (-1G) targets, rendered on two distinct quasi-realistic visual scenarios (Figure 1D and 1E). Temporal intervals were cued by two flanking objects moving with same kinematics (motion duration = 750 ms) but shifted temporally so that their alternated bouncing against flat surfaces defined the interval duration of the visual metronome. Subjects were instructed to tap in sync with the target bounces. As control conditions, we used flashing visual metronomes consisting of alternated flashing for 150 ms of two flanking targets (same visual appearance as the moving conditions) at screen positions corresponding to the bouncing location of the moving conditions (Figure 1F and 1G). Five target interval durations were used (*t_d_*, from 450 to 850 ms in steps of 100 ms) across all visual motion and flashing conditions.

**Figure 1.**
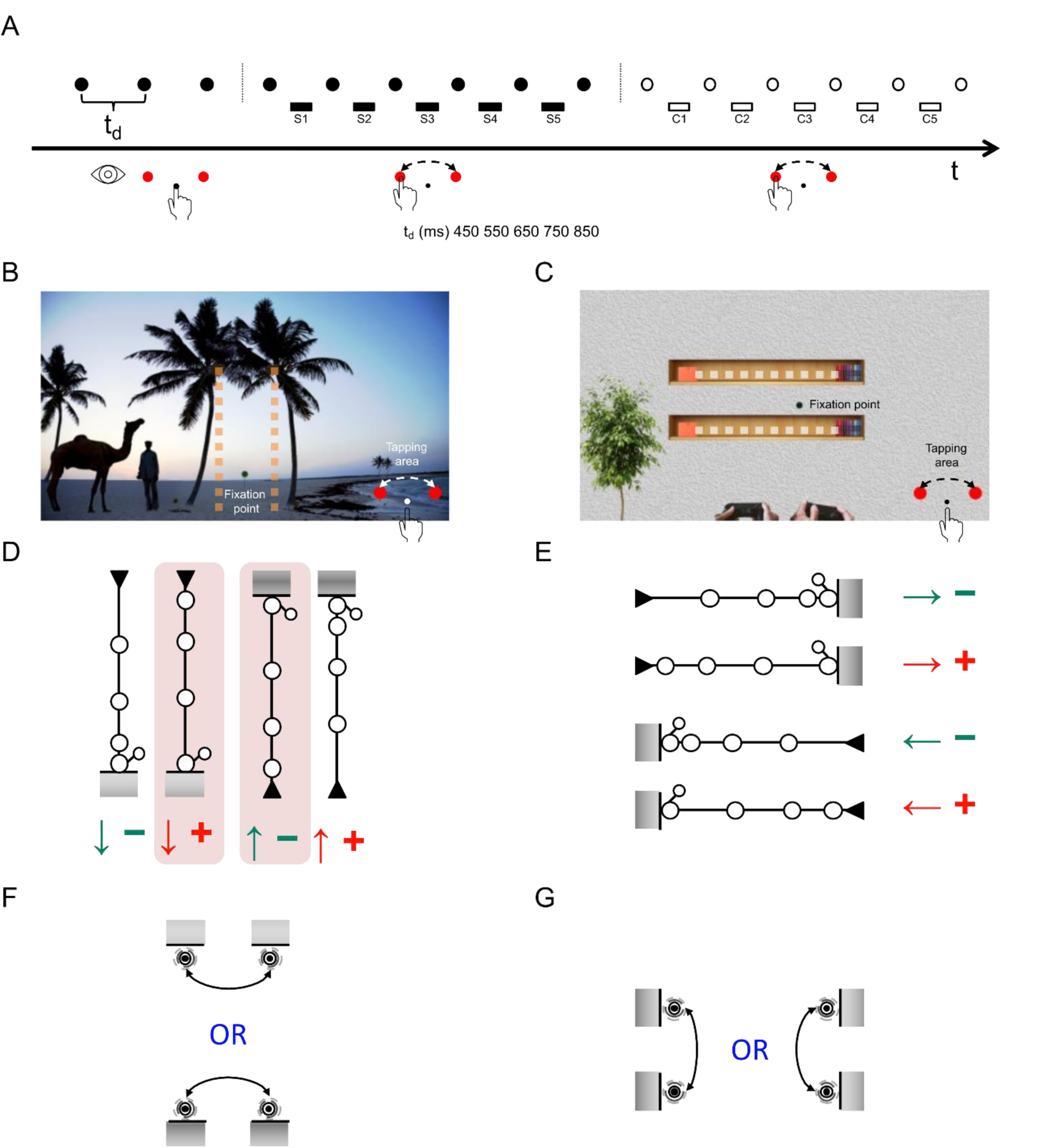
Synchronization continuation task (SCT). (A) Sequence of taps and definition of reproduced temporal intervals. A trial started when subjects placed their right index finger on the starting position at the center of the tapping area located at the right bottom corner of the touchscreen (see also panels B and C). Then, stimuli were presented alternatively in two positions as an isochronous metronome, and subjects, after the first three instruction visual stimuli (observation epoch), produced six taps in synchrony with the metronome (synchronization epoch, *S*1 – *S*5 denote the reproduced intervals), by tapping first on the left red circle and then by alternating between the two red circles in the tapping area. After the visual metronome was extinguished, they continued tapping with the same tempo for other six taps in the absence of any visual cue (continuation epoch, *C*1 – *C*5 denote the reproduced intervals). Five target interval durations were used (*t_d_*, from 450 to 850 ms in steps of 100 ms). During the task, subjects were required to maintain ocular fixation on designated points of the visual scene (see panels B and C). (B) Vertical Scenario. Visual metronomes were presented in separate trials, either at the bottom or at the top of the palm trees, by alternating visual stimuli at fixed intervals between the right (first) and the left tree. The dark green circle between the two palm trees indicates the ocular fixation point when the visual metronomes were presented at the bottom of the trees. The fixation point (not shown for clarity) for visual metronomes presented at the treetops was located between the two palm trees at the same visual angle distance as the one used for the bottom visual metronomes. (C) Horizontal Scenario. Visual metronomes were presented either at the right or left ends of the two bookshelves. The dark green circle between the two shelves denotes the ocular fixation point when visual metronomes were presented at the right end of the shelves. The fixation point for the visual metronomes presented at the left end of the shelves (not shown for clarity) was located between the bookshelves at the same visual angle distance from the visual metronomes at the right end of the shelves. (D) Moving metronomes in the Vertical Scenario. Two coconuts were either dropped from the right and left palm trees and bounced on the ground below the trees or were launched from the ground and bounced against the top branches of the palm trees. In each trial, the two coconuts moved along the same direction and with the same kinematics (motion duration = 750 ms), but their motion onsets were shifted temporally so that alternated bounces between the right (occurring first) and the left coconut occurred at fixed intervals corresponding to one of the 5 possible interval durations of the visual metronome. For each direction of motion, coconuts’ motion could be either accelerated (signified by the open circles going further apart as they approach the bounce, coded as a red plus sign) or decelerated (open circles get progressively closer as they approach the bounce, coded as a minus green sign). Objects’ velocity at the bounce was 59 and 5 visual degrees*sec^-1^ for accelerated and decelerated motion, respectively. After the bounce, coconuts moved at an oblique angle from the bouncing surface and disappeared quickly behind the trees by moving at a velocity comprised between 18 and 21 visual degrees*sec^-1^. The vertical motion conditions congruent with natural gravity effects (i.e. downward accelerated and upward decelerated) are shaded in pink. (E) Moving metronomes in the Horizontal Scenario. Two toy racing cars ran along the upper and lower bookshelves, starting from one end of the bookshelves (symbolized by the filled triangle) and bounced against the pile of books (symbolized by the shaded rectangle) at the opposite end. As in the Vertical scenario, in each trial, the two moving targets had the same kinematics (either accelerated or decelerated; motion duration = 750 ms), but their motion onsets were shifted temporally so that the alternated bounces between the car on the lower (occurring first) and on the upper shelf defined one of the 5 possible interval durations of the visual metronome. Objects’ velocity at the bounce was 59 and 5 visual degrees*sec^-1^ for accelerated and decelerated motion, respectively. After the bounce, cars moved at an oblique angle from the bouncing surface and disappeared quickly behind the book piles by moving at a velocity comprised between 18 and 21 visual degrees*sec^-1^. (F) Flashing metronomes in the Vertical Scenario. In each trial, static coconuts’ images were flashed for 150 ms, alternatively between the right (first) and the left palm tree, defining one of the 5 possible interval durations of the visual metronome. Alternated flashing of the coconuts occurred at the bouncing locations of the moving metronomes, either at the treetops or on the ground beneath the trees. (G) Flashing metronomes in the Horizontal Scenario. In each trial, static toy cars’ images were flashed for 150 ms, alternatively between the bottom (first) and the top bookshelves, at intervals corresponding to one of the 5 possible target durations of the visual metronome. Alternated flashing of the toy cars occurred at the bouncing locations of the moving metronomes, either at the left or at the right end of the bookshelves.

### Precision, accuracy, and prediction of rhythmic timing

First, we assessed the effects of the various experimental manipulations of the moving and flashing metronomes on the constant error, the temporal variability, and the asynchronies. The constant error is a measure of timing accuracy and corresponds to the difference between the produced and target intervals. The temporal variability is a measure of timing precision and corresponds to the standard deviation of the produced intervals. Asynchronies are the time differences between stimuli and tap responses, and represent measures of rhythmic prediction, which can be evaluated only during the synchronization epoch of SCT (Merchant et al., 2008b; Gámez et al., 2018; García-Saldivar et al., 2022).

### Tapping precision

The slope method is a classical timing model that uses a linear regression between temporal variability as a function of target duration (*t_d_*) to arrive at a generalized form of Weber’s law (Figure 2A, data from the first produced interval of the continuation epoch) (Ivry and Hazeltine, 1995). The resulting slope (*sTV*) is associated with the time-dependent process since it captures the scalar property of interval timing. The intercept (*cTV*) is related to the time-independent component, which is the fraction of variance that remains similar across interval durations and is associated with sensory detection and processing, decision making, memory load, and/or motor execution (Merchant et al., 2008a; Zarco et al., 2009). As a convention we computed the *cTV* at the intermediate target interval of 650 ms instead of at 0ms as usually computed in linear regression (Figure 2A). The *cTV* and *sTV* were computed for each subject and condition though linear regression (Figure S1). Figure 2B shows profile of *sTV* values plotted as a function of *cTV* for the deceleration motion conditions (mean of the 4 directions). It is clearly that *sTV* values plotted as a function of *cTV* follow a U-shape curve across the produced intervals of the synchronization (S1-S5) and the continuation (C1-C5) phases. The time independent component (*cTV*) was large at S1, decreased within the sequence with the smaller value around C1, and showed a rebound at the end of the continuation epoch. The time dependent component showed a systematic decrease during synchronization and a slight increase during continuation. Similar U-shaped curves were observed for the eight motion conditions, with larger *cTV* for the deceleration compared to the acceleration conditions, regardless of the target motion directions (Figure S1B).

**Figure 2.**
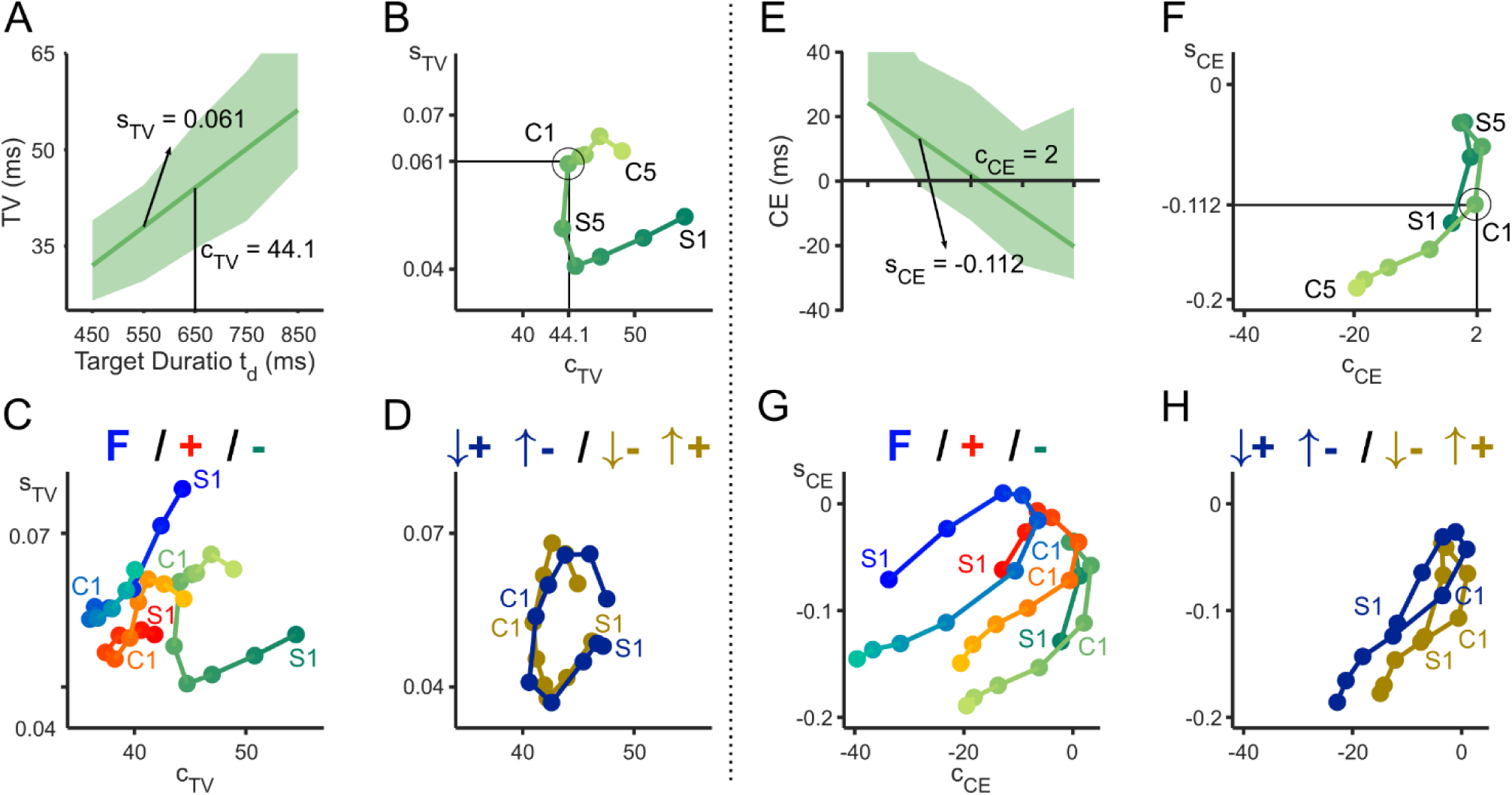
Timing accuracy and precision. (A) The temporal variability (TV; median and interquartiles) of the first interval of the continuation epoch (C1) of the decelerated motion conditions plotted as a function of target duration (*t_d_*). The corresponding linear regression was used to determine the intercept at 650 ms (*cTV =* 44.1 ms), and the slope (*sTV* = 0.061). (B) *sTV* plotted as a function of *cTV* across the serial order elements (S1-S5, C1-C5) of the SCT for the deceleration motion conditions. The circle in C1 highlight the values obtained from A. (C) *sTV* as a function of *cTV* for all serial order elements for the flashing metronomes (blue), all directions with accelerated (red +) and all directions with decelerated (green-) motion. Note the U-shape progression of *sTV* and *cTV* across the serial order elements of the SCT. In addition, there is a large increase in the time independent variability (*cTV*) in deceleration, and an increase in the time dependent variability for the first two serial order elements (S1-S2) of flashing. (D) *sTV* plotted as a function of *cTV* across serial orders by pooling motion conditions in natural and non-natural gravity conditions. Note the overlapping temporal variability profiles in the two gravity conditions. (E) Constant error (CE; median and interquartile) of the first interval of the continuation epoch (C1) with decelerated motion conditions is plotted as a function of *t_d_*. We fitted a linear regression and extracted the intercept at 650 ms (*cCE* = 2 ms), and the slope (*sCE* = −0.112). (F) *sCE* plotted as a function of *cCE* across the produced intervals of the synchronization and continuation for the decelerated motion conditions. The open circle depicts the data from E. Note the inverted U-shape progression of *sCE* and *cCE* across the serial order elements of the SCT. (G) *sCE* as a function of *cCE* for all serial order elements for the flashing metronomes (blue), all directions with accelerated (red +) and all directions with decelerated (green-) motion. Note that *cCE* reached smaller values for flashing, intermediate for acceleration and larger values for the deceleration conditions. (H) *sCE* as a function of *cCE* across serial orders for the natural and non-natural gravity conditions. Note the very similar, but shifted along the abscissa, constant error profiles between the two gravity conditions.

To parametrize these effects, we performed repeated measures ANOVAs using either *cTV* or *sTV* as dependent variable, and the serial order (1-10), stimulus kinematics (acceleration, deceleration), and direction (the four cardinal directions) as factors. The results for *cTV* showed significant main effects of serial order and stimulus kinematics, but neither significant main effects for direction nor significant interactions, except for the significant serial order x stimulus kinematics interaction. In contrast, the *sTV* ANOVA did not show significant main nor interaction effects (Table S1A and Figure S1B). The corresponding repeated measures ANOVAs on the flashing metronome conditions showed only significant main effects of serial order on both *cTV* or *sTV* (Table S1B and Figure S1A). In Figure 2C we compared either *cTV* or *sTV* among flashing, acceleration, and deceleration conditions. It is evident that the responses to decelerated motion show the largest *cTV*, with responses to accelerated and flashing metronomes showing comparable *cTV* values across all the produced intervals in the SCT sequence. Indeed, significant differences emerged only between flashing and decelerated motion (Table S1C and Figure 2C). In addition, *cTV* and *sTV* values in the first synchronization interval (S1) with flashing metronomes are quite larger and significantly different than those observed with moving metronomes (*cTV*: F(2,31) = 27.5, p < 0.0001; all paired t-tests p < 0.0001; *sTV*: F(2,31) = 6.5, p < 0.0001; flashing against acceleration and deceleration conditions with paired t-tests p < 0.05). Crucially, we also tested the effects of natural and non-natural gravity conditions in a separate repeated measures ANOVA, with no significant differences for both *cTV* and *sTV* (Table S1D and Figure 2D).

These results suggest that visual motion produced changes mostly on the time-independent component of the SCT, with lower variability in response to accelerated motion and flashing, and highest for decelerated motion. With respect to the serial order in rhythmic production, larger and smaller variability in both time independent and dependent components was observed for S1 and S5, respectively. No effect of visual motion naturalness was found.

### Tapping accuracy

Figure 2E shows the constant error as a function of *t_d_* for the first produced interval of the continuation epoch of the decelerated motion conditions, with the typical bias effect of overestimation for short and underestimation for long intervals. We defined two measures to characterize this behavior using a linear regression on this data: the intercept at 650 ms (*cCE*), and the slope of the regression called *sCE* (Figure 2E). The former is proportional to the indifference interval, namely, the interval where there is no error in timing (McAuley and Jones, 2003; Rajendran et al., 2018; García-Saldivar et al., 2022). The latter corresponds to the magnitude of the bias effect, i.e., the larger the negative slope, the larger the regression towards the mean (Jazayeri and Shadlen, 2010; Petzschner et al., 2015; Perez and Merchant, 2018). In Figure 2F we plotted profile of *sCE* as a function of the *cCE* for each produced interval in the SCT sequence for the deceleration motion conditions. It is evident that there is an inverted U-shape in the change of these two parameters across the SCT sequence for all motion conditions. At the beginning of the task (S1), there was a negative *cCE* that means an overestimation of *t_p_*, then *cCE* migrating towards cero in the next synchronization intervals (S2-S5) suggesting that indifference interval is around 650ms. During continuation (C1-C5), *cCE* became again large and negative and acquired stationary values. These results indicate three crucial effects of the sequence on timing accuracy: (1) the initial interval in the rhythmic sequence shows a large bias effect, (2) during the next synchronization intervals the accuracy is close to perfect, and (3) during the continuation phase the bias effect resumes with a stable *cCE* that may reflect the subject’s preferred interval under internal rhythmic tapping. Similar trends were observed for all motion conditions, as well as for the flashing conditions (Figure S1C and S1D). Repeated measures ANOVAs showed, for the moving metronomes conditions, significant main effects of serial order and motion kinematics, but neither significant main effects for direction nor significant interactions for both *sCE* and *cCE* (Table S1A). Moreover, for the flashing conditions, the repeated measures ANOVA showed only significant main effects for serial order on *sCE* and *cCE* (Table S1B). We compared the *sCE* vs. *cCE* patterns across serial orders between flashing, acceleration, and deceleration conditions. The inverted U shape in the serial order was the prevalent pattern across all conditions (Figure 2G), however, with a significant increasing shift in the *cCE* profile between flashing, acceleration, and deceleration conditions (all paired t-tests showing *p <* 0.0001, see Table S1C). In addition, the *sCE*, which is a measure of the bias effect, was also larger in the deceleration than in the flashing and acceleration conditions (Table S1C). Crucially, statistical differences were found between natural and non-natural gravity conditions for *cCE* but not for *sCE* (Figure 2H and Table S1D).

These findings indicate that *cCE* was shorter for flashing and larger for deceleration. Although there was a strong tendency to reach an indifferent interval around 550 ms at the end of the continuation epoch for all motion conditions (Figure S2), the overall *cCE* was significantly larger in the non-natural than the natural condition. Finally, the bias effect was larger for deceleration than acceleration and flashing conditions.

### Tapping prediction

The asynchronies showed large negative values with decelerated motion and similar values with flashing and accelerated motion, which were closer to zero, but with a systematic decreasing trend as a function of *t_d_* and of serial order (Figure 3A *top*). Indeed, the repeated measures ANOVA applied to the moving conditions showed significant main effects of motion kinematics and *t_d_*, as well as significant interactions for motion kinematics x *t_d_*, motion kinematics x serial order and serial order x *t_d_* (Table S2A). With flashing metronomes, asynchronies were influenced significantly only by *t_d_* and serial order (Blue traces in Figure 3A *top* and Table S2B). By comparing mean asynchronies with the Static-Kinetic model (Table S2C), we did find significant differences between deceleration and either acceleration or flashing conditions, while asynchronies in response to accelerated and flashing stimuli were statistically indistinguishable (compare red and blue traces in Figure 3A top). Visual motion naturalness did not have statistically significant effects (Figure 3A *bottom* and Table S2D).

**Figure 3.**
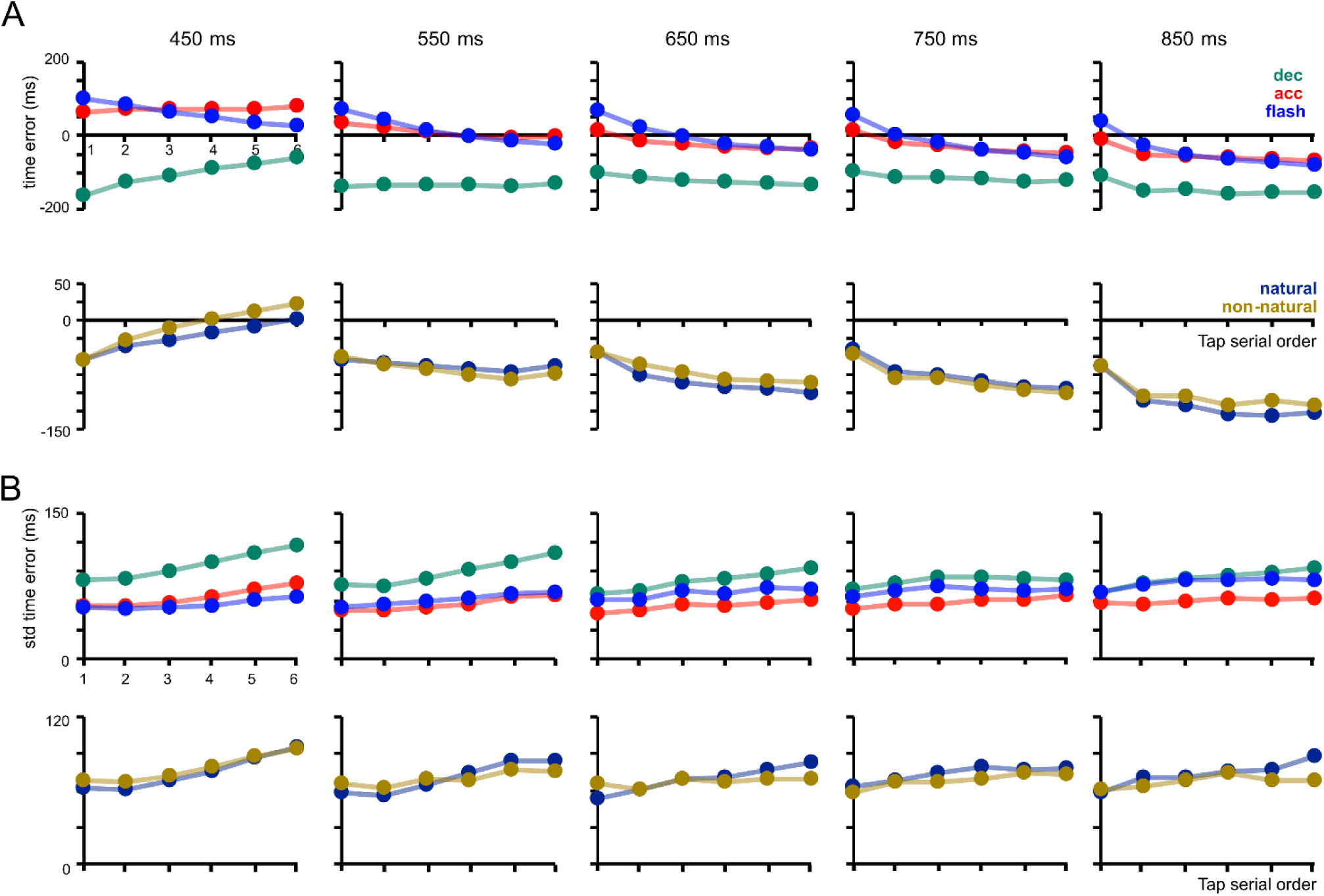
Tapping asynchronies. (A) Top row: tapping asynchronies in response to accelerated (red), decelerated (green) and flashing (blue) visual stimuli. Each graph shows the mean asynchrony values (± SD) computed across subjects, for the series of the six taps in the synchronization epoch for each of the five *t_d_* (label at the top). Bottom row: tapping asynchronies for the vertical motion conditions congruent with natural (blue) and non-natural (brown) gravity. Each graph shows the mean asynchrony values (± SD) computed across subjects, for the six taps of the synchronization phase for each *t_d_*. (B) Top row: Variability in the asynchronies computed as Standard Deviations across trials as a function of the serial order of taps for the same experimental conditions in A top. Bottom row: Variability in the asynchronies for the conditions in A bottom.

The variability of the asynchronies, which corresponds to their intra-trial standard deviation, were significantly larger for the deceleration than the acceleration condition (especially for longer *t_d_*), and showed a systematic increase with serial order, accounted for by a significant serial order x *t_d_* interaction (Figure 3B *top* and Table S2A). With flashing visual metronomes, tap variability increased significantly with the tap serial order and with the *t_d_* (Table S2B). Notably, the variability of the asynchronies was significantly lower not only by comparing acceleration with deceleration conditions, but also between acceleration and flashing stimuli, indicating more precise prediction for the acceleration condition (Table S2C). Finally, we found a small interaction effect between the naturalness of the visual motion and the serial order of taps (Figure 3B *bottom* and Table S2D).

In sum, the results of these analyses indicated that: 1) with both moving and flashing metronomes, the initial interval in the rhythmic sequence showed a large bias effect and a larger time independent component, but both accuracy and precision improved throughout the synchronization phase. The time dependent and independent components as well as the bias effect increased again during the continuation phase, with stable cCE values at the end of continuation phase reflecting the subject’s internalized preferred interval; 2) with moving metronomes, the acceleration of the moving objects was the strongest factor determining the precision and the accuracy of the SCT performance, whereas the effects of the naturalness of motion were limited to the increase of the indifference interval; 3) compared to flashing metronomes, moving metronomes provided a significant advantage in SCT performance only when time intervals were cued by accelerating objects.

### A Bayesian model for rhythmic tapping

In order to gain further insight on the dynamic evolution of the subjects’ SCT performance across the different experimental conditions, we adapted the three stage (measurement, estimate and production) Bayesian model designed for single interval production by Jazayeri and Shadlen (Jazayeri and Shadlen, 2010) to our SCT (Figure 4; see Methods). Initially, the model incorporates the change from an open (interval S1) to a close-loop (intervals S2-S5 and C1-C5) in timing production (Figure 4A). Typically, during the synchronization epoch, an error correction mechanism generates the tendency for short-produced intervals to be followed by a long interval and vice versa, thereby avoiding large error accumulation that could drive tapping responses out of sync with the metronome. In contrast, during the continuation epoch, subjects use an internal clock that tends to generate a small drift in the produced intervals towards progressively shorter or longer intervals. Hence, to capture both phenomena we added a fourth stage to the model (Figure 4B). In this step, the difference between the previously produced interval (*t_p_*_−1_) and the physical interval (*t_d_*) influences the input time as *t_s_* = *t_d_* + *λ* (*t_p_*_−1_ − *t_d_*) with a λ weight. On the first produced interval λ*=* 0 (Figure 4C, λ middle). On the following intervals of the synchronization λ depends on how large error correction adjusts the actual *t_p_*, with values below zero since there is a negative feedback loop (Figure 4C, *λ* top) (see the lag-1 autocorrelation between consecutive intervals in the SCT in Figure S3A). Finally, during the continuation *λ* tends to be positive due to the positive loop of produced intervals causing a drift (Figure 4C, *λ* bottom; see also the drift of produced intervals during the continuation epoch in Figure S3B).

**Figure 4.**
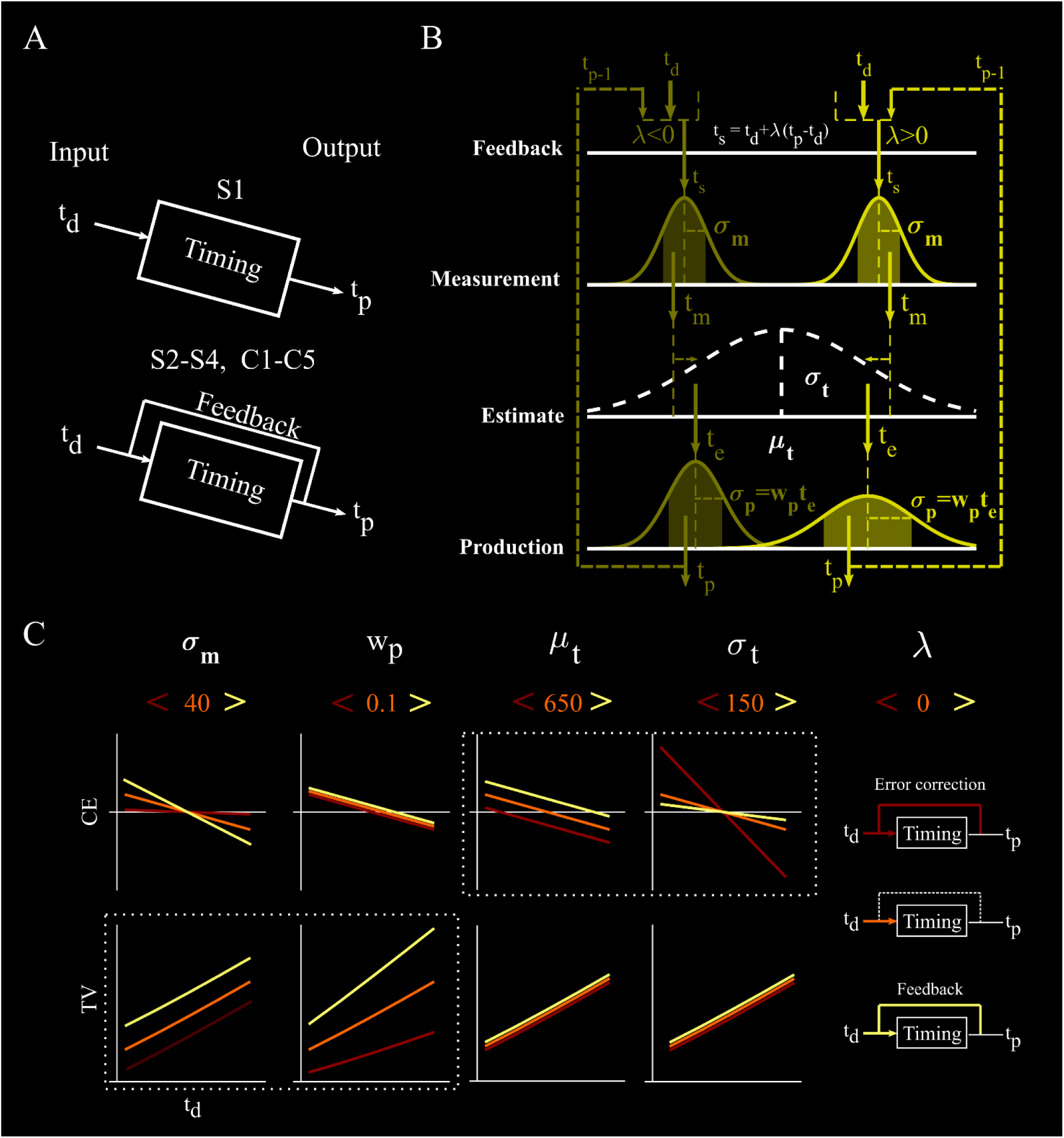
Bayesian model. (A) The model incorporates the change from an open- (top, initial interval S1) to a close-loop (bottom, S2-S5 and C1—C5) in timing production. During the close loop the previous interval has an influence on the current duration allowing for either error correction (negative feedback) or interval drift (positive feedback). (B) The model has a four-stage architecture. First, the target duration *t_d_* defined by the metronome is measured with *t_m_* that is modelled with a Gaussian distribution centered in *t_s_* and that adds noise *σ_m_* on the relation *p*(*t_m_*|*t_s_*). Next, the observer computes a time estimate (*t_e_*) that is the maximum of the posterior probability that is proportional to the product of the likelihood function *p*(*t_m_*|*t_s_*) and the prior probability distribution *p*(*t_s_*) (a Gaussian with a mean *μ_t_*, and a standard deviation *σ_t_*). Third, *t_e_* is used to compute the produced interval *t_p_*, with a conditional dependence of *t_p_* on *t_e_*, *p*(*t_p_*|*t_e_*), which adds noise to the production phase by using Gaussian distribution whose standard deviation is *w_p_t_e_*. The *w_p_t_e_* is larger for longer estimated intervals, simulating the scalar property of timing. The last stage considers the influence of the previous *t_p_*_−1_ on the current interval, computing the difference between the previously produced and the target duration as *t_s_* = *t_d_* + *λ*(*t_p_*_−1_ − *t_d_*). This difference influences the current measured time with a *λ* weight. Thus, on the first produced interval *λ* = 0, but on the following intervals the impact of the previous produced duration depends on how large the negative feedback error correction or the positive feedback drift adjusts the actual *t_p_*. (C) Simulations of the effect of the small (red), intermediate (orange), and large (yellow) values of the parameters of the model on the Constant Error (CE) and Temporal variability (TV). The increase in *σ_m_* produces large changes in the time independent variability (cTV) and a moderate increase in the bias effect. The increase in *w_p_* induces a large increment in cTV and sTV with no changes in CE. An increase in *μ_t_* produces a shift towards the right in the II and no changes in TV. A rise in *σ_t_* induces an increase in the bias effect. Finally, *λ* = 0 implies no feedback, *λ* < 0 involves negative feedback error correction, and *λ >* 0 entail positive feedback with interval drift.

It is important to consider that the noise parameters *σ_m_* for time measurement and *w_p_* for time production, capture the changes in temporal variability during the SCT. In contrast, *μ_t_* and *σ_t_* capture the effects on the constant error, with the former linked to the *cCE* and the indifference interval and the latter related to the magnitude of the bias effect (Figure 4C).

For each subject, we fitted the parameters on the model, namely, *σ_m_*, *μ_t_*, *σ_t_*, *w_p_*, and *λ* on the SCT for the flashing and the eight visual motion conditions. The values of goodness-of-fit were large, with significant correlations between the predicted and actual data across subjects (Figure S4A and S4B). Figure 5 shows the coefficients of the model across the serial orders of the eight motion and the flashing conditions. A repeated measures ANOVA applied to the parameters fitted to the motion conditions indicated that the time measurement noise (*σ_m_*) showed significant main effects of serial order, motion kinematics and their interaction (Figure 5A top and Table S3A), with larger values for S1 across kinematic conditions and smaller values for the acceleration than the deceleration conditions. *σ_m_* also showed significant main effects of serial order and metronome position in the ANOVA applied to the flashing conditions (Table S3B). The static-kinetic ANOVA showed significant effects, with pair-wise differences between deceleration and either flashing or acceleration conditions (Figure 5A *bottom* and Table S3C). The time production noise (*w_p_*), which is associated with the slope of the scalar property, showed a significant main effect of serial order and kinematics (Figure 5B *top* and Table S3A) for the moving metronome conditions and only a main effect of serial order with flashing metronomes (Table S3B). Larger *w_p_* were observed for S1 (Figure 5B *bottom*) and a significantly larger *w_p_* in the flashing than in the accelerated motion condition (Table S3C). The mean of the prior distribution (*μ_t_*), a parameter linked to the indifference interval and *cCE*, showed significant main effects of serial order and motion kinematics for the Moving metronomes ANOVA, and only of serial order for the Flashing metronomes ANOVA (Figure 5C *top* and Table S3A, S3B). It was larger for deceleration than acceleration and flashing (Table S3C), and larger for the synchronization than the continuation epoch (Figure 5C *bottom*).

**Figure 5.**
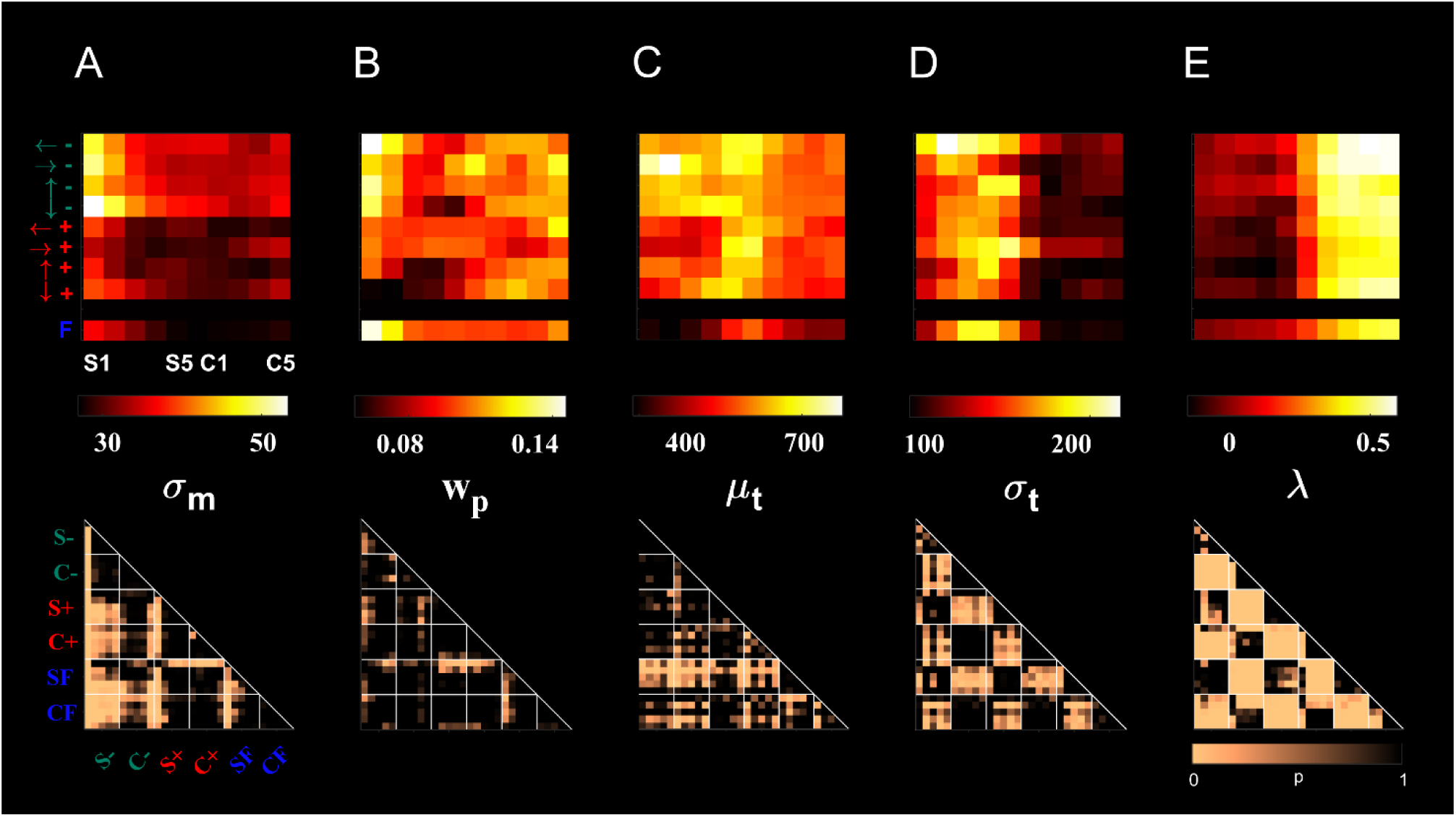
Parameters of the Bayesian model across the flashing and motion conditions. (A) Top: *σ_m_* is plotted for the eight motion (arrows specify the direction, green for deceleration, red for acceleration) and the flashing (Blue F) conditions as a function of serial order in the SCT ranging from S1 to C5. The scale on the heatmap is at the bottom. Bottom: Diagonal matrix of the pair-wise Bonferroni post-hoc test between the five serial order elements of the synchronization and the corresponding five of the continuation epochs, for the four deceleration (green), four acceleration (red), and the flashing conditions (blue F). (B, C, D, E) Values of *μ_t_*, *σ_t_*, *w_p_*, and *λ*, respectively, plotted as in A.

The sigma of the prior (*σ_t_*), a parameter associated with the magnitude of the bias effect, showed a significant main effect of serial order for both the Motion and Flashing metronomes the bias effect was also more evident (Figure 5D). *σ_t_* was larger for deceleration than acceleration and flashing (Table S3C). Finally, *λ*, a parameter that captures the correlation between consecutively produced intervals in the SCT, showed significant main effects of serial order and motion kinematics for the Moving metronomes ANOVA, and only of serial order for the Flashing metronomes ANOVA (Figure 5E *top* and Tables S3A, S3B). *λ* was negative only for the synchronization epoch of the accelerated motion conditions and significantly different from the decelerated ones, indicating a robust error correction (Table S3C). Moreover, *λ* was positive and larger for the continuation than the synchronization epoch (Figure 5E *bottom*) capturing the drift in produced intervals during the internally driven phase of the SCT.

Overall, the Bayesian Model captured all the precision and accuracy properties of rhythmic performance on the SCT across target conditions described above (Figure S4C). Crucially, it also indicated that, independently of the target condition, subjects optimally switched between the sensory guided tap synchronization and the internally driven continuation, by using mechanisms of error correction and regression towards the mean, respectively. It is noteworthy that none of the Bayesian Model parameters, except for *μ_t_*, showed significant differences between the natural and non-natural gravity conditions (Table S3D).

## Discussion

The results of the present study support three main conclusions. First, when syncing to moving metronomes, timing accuracy, precision, and the correlation of consecutively produced intervals were more efficient with accelerated than decelerated motion and hardly influenced by the object motion naturalness, implying that motion kinematics per se was the predominant factor determining rhythm synchronization. Second, metronomes cued by accelerated motion provided an overall performance advantage relative to flashing visual metronomes. Third, a Bayesian observer model could account for the changes in the bias effect, the scalar property, as well as the error correction or drift in produced intervals within the synchronization and continuation epochs of the SCT across motion conditions, providing a parametric account of isochronic rhythmic behavior that indicates an optimal shift of behavior depending on the epoch of the SCT.

### Effects of motion kinematics and naturalness on rhythmic timing

It has been previously shown that the auditory–visual asymmetry for beat induction can be canceled out by using visual moving metronomes (Hove et al., 2010), especially with naturalistic stimuli, such as videos of a continuous bouncing ball with changes in speed (Hove et al., 2013; Gan et al., 2015; Iversen et al., 2015). One hypothesis to explain these observations is that visual motion may engage the time to collision mechanism underlying the control of interceptive actions or of collision avoidance (Merchant et al., 2004a; Merchant and Georgopoulos, 2006). According to this idea, the parieto-premotor system recruited for encoding time-to-contact for discrete events could also be involved in the internal representation of periodic collision points, driving efficiently the motor system for beat-based timing (Mendoza and Merchant, 2014; Iversen et al., 2015; Comstock et al., 2018). The present study was designed to test whether physically realistic motion trajectories with downward-acceleration or upward-deceleration profiles induced better rhythm entraining in the SCT than the non-naturalistic upward-acceleration/downward-deceleration or accelerating/decelerating horizontal motion metronomes. In effect, a natural gravity gain on rhythmic timing might be expected based on the evidence that diving gannets use gravitational signals to compute the time-to-contact and fold their wings before entering the water (Lee and Reddish, 1981), and that humans can use an internal representation of gravity effects residing in the vestibular cortex to time accurately the interception of vertically falling objects (Lacquaniti and Maioli, 1989; Indovina et al., 2005; Lacquaniti et al., 2015; Delle Monache et al., 2021). However, our findings were not congruent with this expectation, since the timing accuracy, precision, and the correlation of consecutively produced intervals were not different between the natural and non-natural gravity vertical motion nor with horizontal motion. Rather, they suggest that the parieto-premotor internal representation of periodic collision points either does not integrate or down weights signals from the parieto-insular vestibular cortex – which are known to encode a-priori information about gravity (Indovina et al., 2005) – to generate the beat representation during SCT. The only consistent effect of the naturalness of the visual motion was a shortening of the indifference interval, implying that gravity related information may tune the preferred tempo representation.

Our results indicated also that the parieto-premotor system entrains more efficiently rhythmic tapping with metronomes cued by accelerated motion compared to decelerated motion. Neurons in middle temporal (MT) extrastriate visual cortex, which is reciprocally connected with the posterior parietal cortex (Cavada and Goldman-Rakic, 1993), are tuned for direction and speed of moving visual stimuli but not to acceleration-deceleration (Movshon et al., 1990; Lisberger and Movshon, 1999). However, recent studies have shown that acceleration information can be extracted by assessing changes over time in the stimulus speed population code in MT (Price et al., 2005; Schlack and Albright, 2007).

We could, then, hypothesize that the reader of speed population signals from MT in the parietal-premotor system could build a stronger rhythmic timing prediction with accelerating compared to decelerating motion. For example, this could be achieved by a better estimation of the difference between the initial object velocity and the velocity at the bounce in the accelerated condition, due to a bias in the reading system towards increasing speeds. In other words, the stronger internal representation of periodic collision points may depend on the more robust estimation of motion acceleration. In addition, the higher terminal velocity and the sharper speed change across the bounce occurring with accelerating objects, by producing higher and sharper levels of activity in MT neurons tuned to high speeds, could have entrained sharper and less variable ramping activities in parietal and premotor neurons encoding the event timing and controlling the rhythmic tapping behavior (Merchant et al., 2011; Merchant and Averbeck, 2017).

### Preferred tempo and the internally produced intervals

Classical tapping studies coined the notion of preferred rhythmic tempo, which corresponds to the interval produced naturally when asked to tap in the absence of external cues (Fraisse, 1963, 1978; McAuley et al., 2006). The preferred tempo in human adults is around 600ms (García-Saldivar et al., 2022). The present findings indicate that, during the last three intervals of the continuation epoch, the subjects’ indifference interval converged to a particular value that was shorter for flashing than accelerating and decelerating metronomes and was larger for non-natural than natural gravity motion (Figure 2). These findings suggest that the internal timing mechanism use the bias effect to cope with the lack of entraining sensory input across conditions, but the preferred tempo is directly affected by motion properties of the metronome, changing the indifference interval even during continuation where the stimuli were turned off several intervals before.

### Bayesian modelling of rhythmic timing

The results of the model pointed out several interesting features pertaining the synchronization strategies adopted with moving and flashing metronomes: a) Subjects used three different optimal strategies to produce intervals within the tapping sequence: a large mix between prior knowledge of the statistics of used intervals and the time measurement at the beginning of the synchronization epoch, which is very sensitive to the properties of the metronome; b) an error correction mechanism during the successive intervals of the synchronization that compensated for the effect of the prior and increased the timing precision; c) a balanced use of prior and time measurement during the internally driven epoch of the task, with a large feedforward influence between consecutive intervals producing the characteristic drift of the continuation; d) the indifference interval was modulated by serial order but reached homogenous values at the end of the continuation, suggesting the expression of a preferred interval that was modulated by the metronomés properties.

The use of Bayesian inference to describe timing performance has been widely employed to characterize the optimal combination of the prior knowledge about the statistics of the presented intervals of an experiment with the likelihood function of the measured time (Petzschner et al., 2015). This modeling approach offers some advantages over more classical approaches such as the iconic Wing and Kristofferson model (Wing and Kristofferson, 1973a, 1973b), which focused only on the continuation epoch and was designed to split the total variability into a motor and a scalar timing component, neither including the timing accuracy nor the change in correlation between consecutively produced intervals in the two epochs of the SCT. Bayesian models, like the one we adopted, not only can capture the scalar property of timing by adding noise that is proportional to the interval during time production, but can also explain the bias, as well as the negative or positive correlation of lag 1 intervals. Particularly, in the Bayesian framework, the prior represents the bias or regression towards the mean effect, and it has been suggested to be the result of an error minimization strategy (Jazayeri and Shadlen, 2010; Petzschner and Glasauer, 2011; Cicchini et al., 2012). In contrast, our model explains the bias and scalar properties of timing, as well as the negative or positive correlation of lag 1 intervals. Our results indicate that this error optimization strategy was used at the beginning of the SCT sequence and during the internally driven epoch of the trial. However, during synchronization, it was replaced by an error correction mechanism that could adjust the duration of consecutive intervals in a negative feedback fashion (short followed by long, long followed by short). This mechanism allows for the compensation of movement timing to avoid large error accumulation (Repp and Penel, 2004; Jantzen et al., 2017). Previous psychophysical studies in humans have reported strong error correction for auditory and weak error correction for flashing visual metronomes (Iversen et al., 2015; Comstock et al., 2018). Here we found that accelerated but not decelerated motion produced a robust error correction, comparable if not superior to that observed with flashing metronomes (Figures 5E and S3A). Crucially, the parameter lambda that we included in the new version of the Bayesian model by Jazayeri and Shadlen (Jazayeri and Shadlen, 2010; Betancourt et al., 2022) accounted not only for the error correction strategy during synchronization but also for the drift in the duration of the produced intervals during the continuation epoch. These findings support the notion that during rhythmic timing humans use dynamically prior knowledge or error correction to optimize time interval production depending on the task contingencies, adding a new dimension on optimal control of rhythmic behavior.

### Visual timing in practice: kinematics of the conductor’s baton

A real-world example of visual rhythmic timing is in an orchestra, where a conductor, among a range of other expressive features, uses visual gestures to communicate very clear timing cues that enable individuals in an ensemble to produce highly synchronized sound (Globerson et al., 2021). In addition to gesturing beats, the conducting pattern typically follows standard shapes depending on the musical meter, as the 2, 3, and 4-beat hand arm trajectories. Notably, the shapes produced by the conductor are not solely in the vertical plane. Rather, they involve movements in different directions to produce the stereotypical shapes, which cue the musicians’ ensemble. Since each shape represents an entire bar of music consisting of 2, 3 or 4 beats, beats are therefore not conveyed only through vertical movements, which may exploit the effects of gravity. Indeed, empirical studies investigating the kinematics of conducting on synchronization through tapping movements indicated that the acceleration of the baton is the key kinematic variable explaining synchronization behavior (Deliège et al., 2011), more so than the direction of motion, the changes in direction or the instantaneous speed(Luck and Nte, 2008; Luck and Sloboda, 2009). Notwithstanding the differences between the biological motion patterns generated by the orchestra conductor and the inanimate object motion employed in the present study, our finding that accelerating metronomes, regardless of movement direction, produced the most precise and accurate tapping is consistent with these studies of orchestral conducting.

## Conclusion

In sum, our experimental findings and computational modeling link together and add detail to a consistent account of the movement kinematic properties that can induce a visual beat in both laboratory and real-world settings, as well as providing insight on the neural dynamics in parieto-premotor regions that underlie these abilities. Furthermore, our model provides a parametric account not just for the occurrence of taps, but also for the dynamic production of taps on a beat-by-beat basis, given previous sensory and motor history. In particular, the four stages of our model – measurement, internal estimation, motor production, and feedback – may provide a useful framework for deeper quantitative investigation of synchronization to visual cues provided by a conductor. These insights may also be relevant for computational gesture analysis and interactive tools based on conducting (Toh et al., 2013; Schramm et al., 2015), and for our understanding of whole-body movement dynamics during dance (Burger et al., 2014).

## Materials and Methods

### Subjects

Sixteen volunteers (eight females, age 22.94 years ± 2.82 SD) were enrolled for the study. Experimental procedures were previously approved by the ethical committee of the University of Rome “Tor Vergata” (protocol #: 23/18) and participants signed a written consent form before participating to the experiment. Subjects were either right-handed or ambidextrous (one subject) according to the Edinburgh questionnaire and had normal or corrected-to-normal vision. Participants sat 35 cm in front of a 22 inches touchscreen (Elo Touch ET2201L, Elo TouchSystems, USA) where visual scenes were displayed with 1920×1080 pixel resolution and refresh rate of 60 Hz. Scenarios were created using the Matlab (2017b; Mathworks, USA) Psychtoolbox-3 toolbox (Ivry and Hazeltine, 1995). Tapping responses were acquired via the usb interface of the touchscreen at a sampling frequency of 120 Hz. Eye movements were recorded with an Eyelink 1000 tracker system (Sr Research, Canada), at a sampling frequency of 500 Hz.

### Synchronization-Continuation Task

Subjects performed two variations of the Synchronization-Continuation Task (SCT). In one variation they synchronized finger tapping responses to events produced by moving visual objects (Moving metronome condition). In the second variation, which was used as control condition, participants tapped in sync with the flashing of the same visual objects for 150 ms at fixed screen locations (Flashing metronome condition).

Each trial started when subjects placed their right index finger on the starting position at the center of the tapping area located at the right bottom corner of the touchscreen (see Figure 1A). Then, stimuli were presented alternatively in two positions as an isochronous metronome, and subjects, after the first three instruction visual stimuli (observation epoch), produced six taps in synchrony with the metronome (synchronization epoch), by tapping first on the left red circle and then by alternating between the two red circles in the tapping area. After the visual metronome was extinguished, they continued tapping with the same tempo for other six taps in the absence of any visual cue (continuation epoch). Five target interval durations (*t_d_*) were used, from 450 to 850 ms in steps of 100 ms. During the task, subjects were required to maintain ocular fixation on designated points of the visual scenes (see below).

### Vertical and Horizontal Scenarios

Visual metronomes were displayed in two different visual scenarios, which were designed specifically to represent either vertical or horizontal object motion (Vertical and Horizontal Scenario, respectively). The two scenarios were presented in separate experimental sessions, performed at least 24 hours apart, and their order of presentation was counter-balanced across subjects. In the Vertical Scenario, visual metronomes were rendered over a structured background scene reproducing a tropical beach, in which two palm trees surrounded by other naturalistic graphic elements provided cues for scaling the scene to real world size (Figure 1B).

For the Moving metronome condition, two coconuts (9.4 visual ° apart) were either dropped from the right and left palm trees and bounced on the ground below the trees or they were launched from the ground and bounced against the top branches of the palm trees. For each direction of motion, coconuts’ motion could be either accelerated or decelerated at a magnitude of 9.81 m*s^-2^ scaled to the virtual scene size. Objects’ velocity at the bounce was 59 and 5 visual degrees*sec^-1^ for accelerated and decelerated motion, respectively. After bouncing, coconuts moved at an oblique angle from the bouncing surface and disappeared quickly behind the trees by moving at a velocity comprised between 18 and 21 visual degrees*sec^-1^ (Figure 1D and supplemental videos). In each trial, the two coconuts moved along the same direction and with the same kinematics (motion duration was fixed at 750 ms), but their motion onsets were shifted temporally so that alternated bounces between the right (occurring always first) and the left coconut defined one of the five possible *t_d_* of the visual metronome. Note that, in this scenario, downward accelerated and upward decelerated motion of the coconuts represented conditions compatible with the effects of natural gravity, whereas upward accelerated and downward decelerated motion were conditions incongruent with a natural setting.

In the Flashing metronome condition, the static images of the coconuts were flashed for 150 ms at the bouncing locations of the moving metronome conditions, either on the ground or on the tree branches. In each trial, alternated flashing of the two visual objects started always from the right side and occurred at fixed intervals corresponding to one of the five *t_d_* of the visual metronome (Figure 1F).

In the Horizontal Scenario, the two visual objects were represented by remotely controlled toy racing cars positioned upon two bookshelves, separated vertically by 9.4 visual ° (see Figure 1C). In the Moving metronome condition, the toy cars ran along the upper and lower bookshelves, starting from one end of the bookshelves, and bounced against the pile of books at the opposite side of the bookshelves. Objects’ velocity at the bounce were identical to those described for the vertically moving objects. After the bounce, cars moved obliquely from the bouncing surface at a velocity comprised between 18 and 21 visual degrees*sec^-1^ and disappeared quickly behind the book piles. In separate trials, the toy cars moved either rightward or leftward and their motion could be either accelerated or decelerated at 9.81 m*s^-2^ scaled to the virtual scene size (Figure 1E and supplemental videos). Alike the Vertical scenario, the two cars had the same kinematics (motion duration = 750 ms), but their motion onsets were shifted temporally so that alternated bounces between the cars on the lower (occurring first) and on the upper shelf defined one of the five *t_d_* of the visual metronome.

In the flashing condition, static images of the toy cars were flashed for 150 ms at the bouncing locations of the Moving metronome condition, either at the left or at the right end of the bookshelves. In each trial, alternated flashing of the two visual targets started always from the lower shelf and corresponded to one of the five possible *t_d_* (Figure 1G).

During each SCT trial, subjects maintained ocular fixation on a dark green dot (diameter 1.1 visual °) positioned always ∼ 8 visual ° from the visual metronome (see the examples shown in Figure 1B-C for visual metronomes presented at the bottom of the palm trees in the Vertical Scenario and at the right end of the bookshelves in the Horizontal Scenario). The fixation point color turned red between trials to instruct subjects they could break fixation temporarily.

Each experimental session comprised twenty Moving metronome conditions resulting from factoring two motion directions (UM | DM in the Vertical, RM | LM in the horizontal Scenario), two motion accelerations (dec | acc) and five *t_d_* (450 | 550 | 650 | 750 | 850 ms). Similarly, 10 Flashing metronome conditions were obtained by factoring two flashing locations (top | bottom for the Vertical; right | left for the Horizontal Scenario) and five *t_d_* (450 | 550 | 650 | 750 | 850 ms).

A pseudorandom sequence of eight repetitions of these thirty conditions (240 trials total) was presented in six mini-blocks of forty trials each. The first two mini-blocks involved Moving metronome trials, grouped with respect to the visual objects’ movement direction; the third mini-block included Flashing metronome trials presented at the location of the first Moving metronome mini-block. After a short break (∼ 5 min), subjects performed the remaining two Moving metronome mini-blocks (presented in the same order as the first two mini-blocks) and the final mini-block with Flashing metronomes presented at the location of second Moving metronome mini-block. The order of mini blocks with different objects’ motion directions was counterbalanced across participants.

### Data pre-processing

We evaluated subjects’ performance during the SCT by analyzing their tapping response times, acquired at 120 Hz from the touchscreen via a standard USB interface. The series of twelve taps produced during each trial delimited 11-time intervals. The first five produced intervals (*t_p_*) represented subjects’ reproduction of the target duration (*t_d_*) during the Synchronization phase, whereas the last five corresponded to the reproduction of the memorized *t_d_* during the Continuation phase. The sixth interval, delimited by the last tap of the Synchronization phase and the first of the Continuation phase, was discarded. For each subject we computed the mean constant error (CE) and temporal variability (TV) of each *t_p_* of the Synchronization and Continuation phases across trials of the same experimental condition. The constant error can be defined as *t_p_* − *t_d_* and represents a measure of timing accuracy. The temporal variability is a measure of timing precision and corresponds to the standard deviation of the *t_p_*.

From the response times, we also determined the tap asynchronies, that is, the time differences between the taps and the visual events cueing the subjects’ responses during the synchronization phase (objects’ bouncing or flashing, depending on the type of visual metronome). Negative and positive asynchrony values indicated whether subjects anticipated or followed the cueing visual events. Then, for each subject we computed the mean and the standard deviation values of the asynchronies for each tap of the SCT series across trials of a given experimental condition.

These datasets were screened for trials during which subjects either broke ocular fixation or did not perform the tapping sequence correctly. Following this screening procedure, we discarded 127 out of 3840 trials (3.3%) in which subjects’ ocular fixation deviated more than 2° of visual angle for longer than 200 ms; additional 176 trials were discarded (4.6% of the total) because participants did not complete the series of 12 taps; 263 additional trials (6.9% of the total) were discarded because the *t_p_* were above 3 standard deviations from the mean values observed in that experimental condition. Custom-made scripts implemented in MATLAB (Mathworks, USA) were used for data preprocessing.

### Slope analysis

The slope method is a classical timing model that uses a linear regression between temporal variability (TV) as a function of target duration (*t_d_*) to arrive at a generalized form of Weber’s law (see Figure 2A). The resulting slope (*sTV*) is associated with the time-dependent process, capturing the scalar property of interval timing. The intercept (*cTV*) is related to the time-independent component, which is the fraction of variance that relatively invariant across interval durations and is associated with sensory detection and processing, decision making, memory load, and/or motor execution (Ivry and Hazeltine, 1995; Merchant et al., 2008a; Zarco et al., 2009). As a convention, we computed the *cTV* at the intermediate target interval of 650 ms instead of at 0 ms as usually computed in linear regression (Figure 2A). The *cTV* and *sTV* were computed for each subject and condition though linear regression.

Similarly, we regressed the constant error on the *t_d_* and defined two other measures to characterize further the SCT behavior (see Figure 2E): 1) the intercept at 650 ms (*cCE*), which is proportional to the indifference interval, namely, the interval where there is no error in timing (McAuley and Jones, 2003; Rajendran et al., 2018; García-Saldivar et al., 2022); 2) the slope of the regression line called *sCE*, which corresponds to the magnitude of the bias effect, i.e., the larger the negative slope, the larger the regression towards the mean (Jazayeri and Shadlen, 2010; Petzschner et al., 2015; Perez and Merchant, 2018)

### Bayesian model

In order to gain further insight on the dynamic evolution of the subjects’ SCT performance across the different experimental conditions, we adapted the three stage Bayesian model designed for single interval production by Jazayeri and Shadlen (Jazayeri and Shadlen, 2010) to our SCT. Initially, the model incorporates the change from an open (interval S1) to a close-loop (intervals S2-S5 and C1-C5) in timing production (see Figure 4).

We used as initial input data the target *t_d_* and produced intervals *t_p_*. Our four-step model included: 1) A measurement process characterized by *p*(*t_m_*|*t_s_*) = *N*(*t_m_*|*t_s_*, *b_m_*), where *N*(*t*|*μ*, *σ*) is a normal distribution with mean *μ* and standard deviation *σ*; 2) An estimation process given by *t_e_* = *f*(*t_m|_b_m_*, *μ_t_*, *σ_t_*) as the maximum of the posterior probability, which is proportional to the product of the likelihood function *p*(*t_m_*|*t_s_*) and the prior probability distribution *p*(*t_s_*) = *N*(*t_s_*|*μ_t_*, *σ_t_*); 3) a production process described by *p*(*t_p_*|*t_e_*) = *N*(*t_p_*|*t_e_*, *w_p_t_e_*); and 4) a feedback process given by *t_s_* = *t_d_* + *λ*(*t_p_*_−1_ − *t_d_*). All these processes were summarized in the conditional probability:

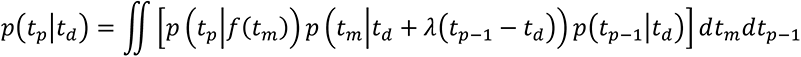

where *b_m_*, *μ_t_*, *σ_t_*, *w_p_* and *λ* were the model parameters (see Figure 4B). Assuming independence of the pairs (*t_d_*, *t_p_*) across different trials, the total conditional probability could be written as 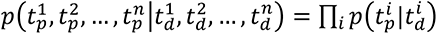. We used this equation to maximize the likelihood probability for the different model parameters, across all pairs of behavioral measures(*t_d_*, *t_p_*). Maximization was carried out with the MATLAB gradient-based method.

### Statistical analyses

The parameters derived from the slope analysis (*sTV*, *cTV*, *sCE*, *cCE*) and from the Bayesian model (*σ_m_*, *μ_t_*, *σ_t_*, *w_p_*, *λ*) were analyzed with four different repeated measures ANOVA models. Two of these models were applied separately to data from either moving or flashing metronome conditions, the third (Static-Kinetic model) tested differences among accelerated decelerated and flashing conditions (thus across the two types of visual metronomes) and the fourth (Gravity model) tested the effects of motion naturalness.

Specifically, for the Moving metronomes model, we included the following “within subject” factors: motion direction (upward | downward | leftward | rightward), motion kinematics (accelerated | decelerated), serial order of *t_p_* (10 levels) and all their two-way interactions.

For the Flashing metronomes model, we considered the following “within subjects” factors: target flashing position (up | down | left | right), serial order of *t_p_* (10 levels) and all their two-way interactions. The Static-Kinetic Model was a one-way repeated measure ANOVA with a three-level factor (accelerated | decelerated | flashing). The Gravity model included data from the vertical session only, collapsed with respect to the naturalness of the vertical motion, and considered the following “within subjects” factors: gravity (natural | non-natural), serial order of *t_p_* (10 levels) and their two-way interaction.

The datasets of mean and standard deviation asynchrony values pooled from the 16 participants were submitted to four repeated measures ANOVA models similar to those described above, with the difference that we considered the serial order of taps during the synchronization phase (6 levels) rather than the serial order of *t_p_*, and that we included the “within subjects” factor target duration (*t_d_*: 450 | 550 | 650 | 750 | 850) to the Moving metronomes, Flashing Metronomes and Gravity models. Greenhouse-Geisser corrections were applied to the ANOVA factors’ p-values with statistical significance cut-off of p = 0.05.

Finally, pairwise comparisons between levels of the ANOVA factors were evaluated by means of two-tailed paired t-tests, Bonferroni corrected (cut-off p = 0.05).

## Acknowledgments

Hugo Merchant is supported by Consejo Nacional de Ciencia y Tecnología (CONACYT): A1-S-8430, UNAM-DGAPA-PAPIIT: IN201721, and SECITI: 2342 to H. Merchant.

Gianfranco Bosco is supported the Italian Ministry of University and Research (PRIN 2017KZNZLN_003) and by the Italian Ministry of Health (RF-2019-12369194).

We are very grateful for the valuable comments that Vani Rajendran provided to our manuscript. We also thank Raul Paulín and Luis Prado for their technical assistance.

---

**Figure S1.**
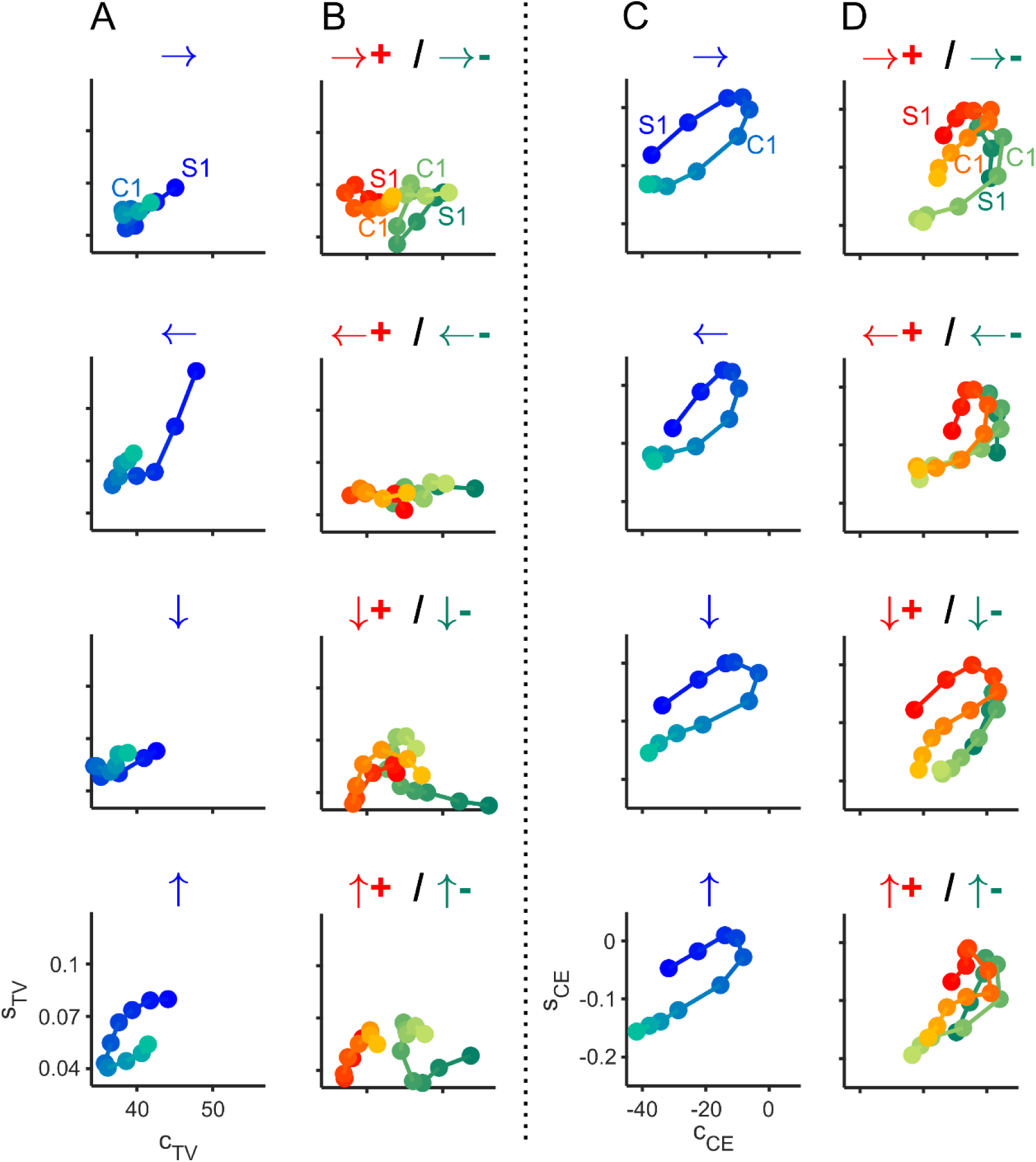
Timing accuracy and precision across all experimental conditions. (A) *sTV* plotted as a function of *cTV* across the serial order elements (S1-S5, C1-C5) of the SCT for the four locations of the flashing stimuli. The top arrow on each of the four panels depicts the position of the flashing metronome in the computer screen. (B) *sTV* plotted against *cTV* for all serial orders of the SCT of the motion conditions. The top arrows on each of the four panels correspond to the motion direction and the acceleration (red, +) and deceleration (green, -) kinetics. (C) *sCE* plotted as a function of *cCE* across the produced intervals of the synchronization and continuation for the flashing metronome conditions. Same notation as in A. (D) *sCE* against *cCE* are plotted for the serial order elements for all motion conditions. Same notation as in B.

**Figure S2.**
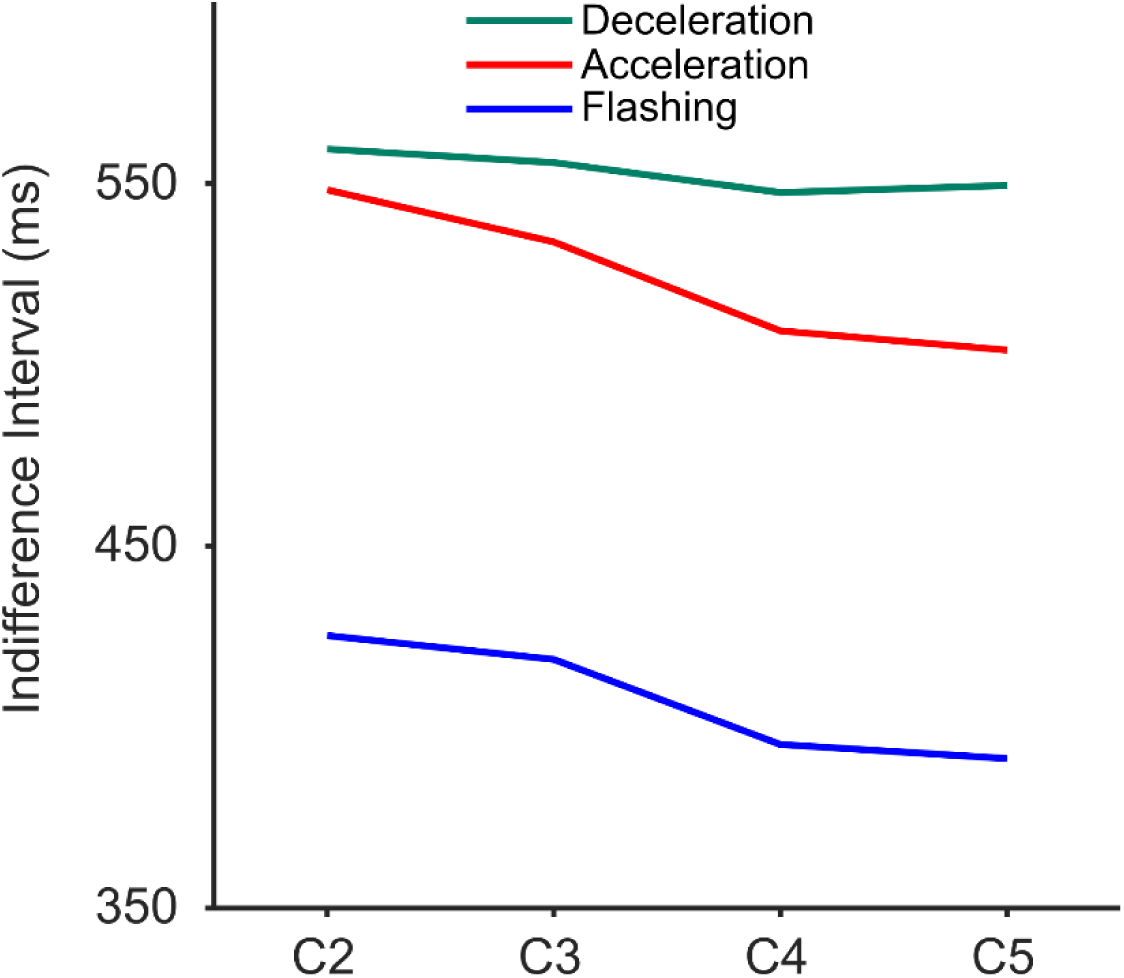
Indifferent interval. Indifferent interval 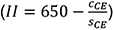 as a function of the last for intervals of the continuation epoch (C2-C5) for the flashing, acceleration, and deceleration conditions (see inset for color code). The static-kinetic ANOVA showed statistically significant differences (F2 = 25.11, p < 0.0001) and the paired t-tests showed differences between flashing and acceleration/ deceleration conditions (p < 0.0001), but not differences between acceleration and deceleration conditions (p = 0.1).

**Figure S3.**
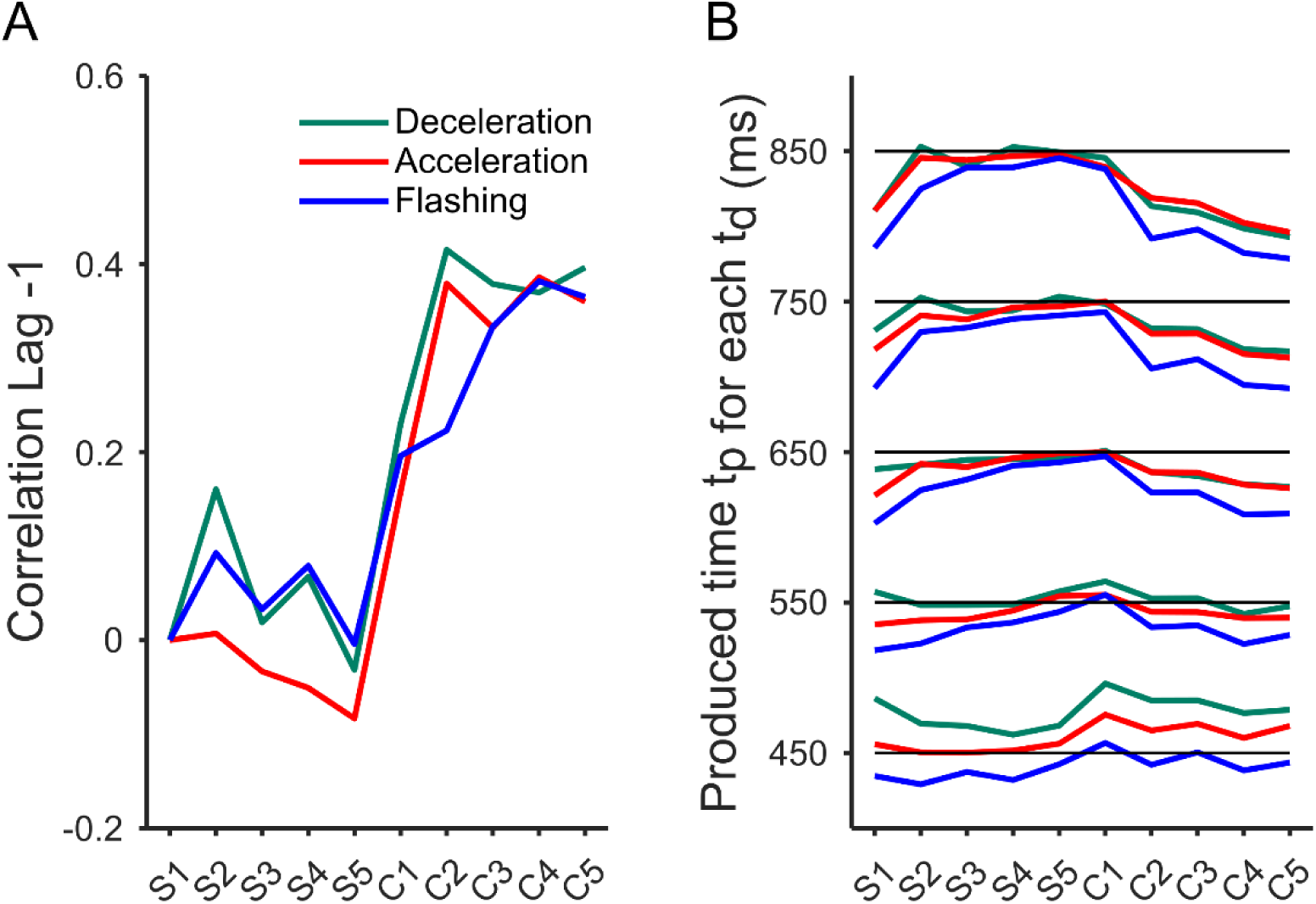
Correlation Lag-1 and Duration of the produced interval. (A) Correlation Lag -1 of the consecutively produced intervals in the five synchronization and five continuation elements for the flashing, acceleration, and deceleration conditions. (B) Produced interval as a function of serial order of the synchronization and continuation epochs for the flashing, acceleration, and deceleration conditions. Important drifts in the continuation epoch are evident for the longer target durations (*t_d_*).

**Figure S4.**
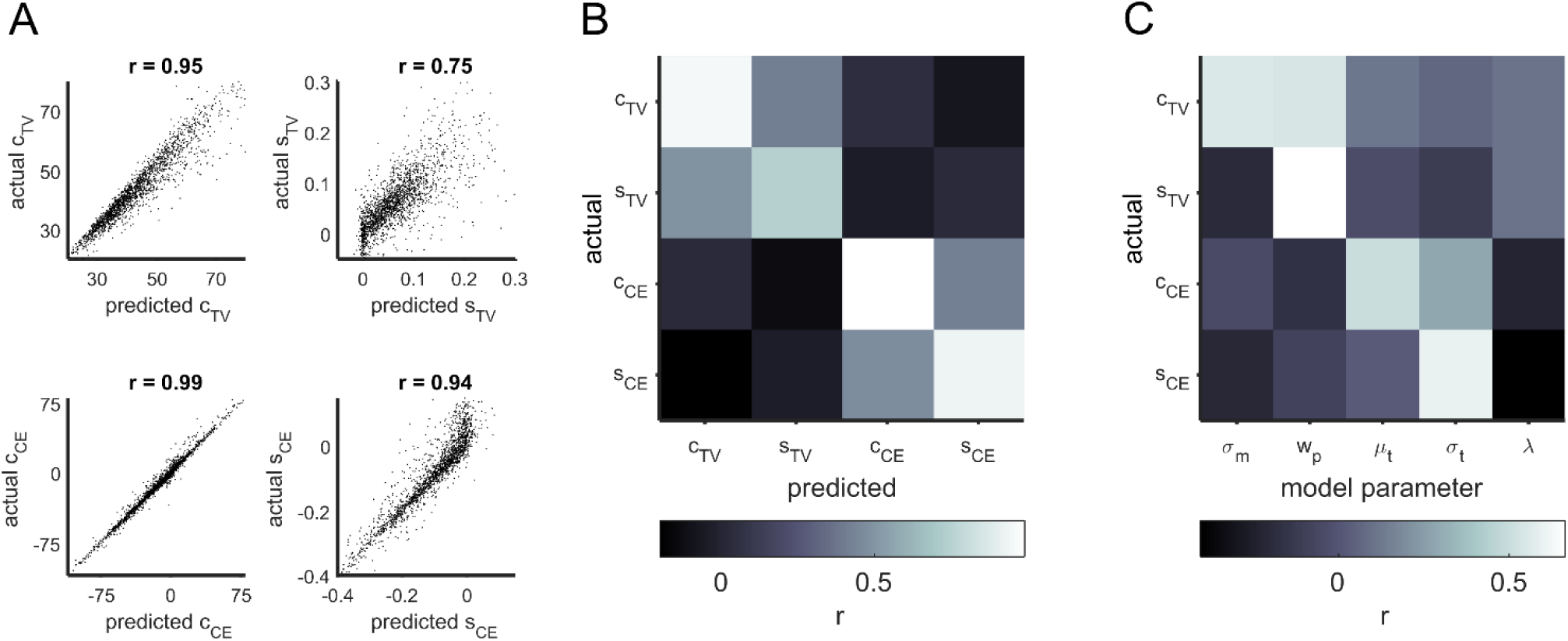
Actual and predicted values of the Bayesian model. (A) Actual against the predicted data for the *cTV*, *sTV*, *cCE* and *sCE* inferred from the Bayesian Model parameters. The significant Pearson correlation values are on top. (B) Pairwise correlation matrix between the actual and predicted data for *cTV*, *sTV*, *cCE* and *sCE* inferred from the Bayesian Model parameters. The inferior bar depicts the color code of the Pearson correlation r value. (C) Pairwise correlation matrix between the actual data for *cTV*, *sTV*, *cCE* and *sCE* and the Bayesian Model parameters.

**Table 1.** Results of the four repeated measures ANOVA models (A-D) applied to the intercept of the Temporal Variability (cTV; time independent component), the slope of the Temporal Variability (sTV; time dependent component), the intercept of the Constant Error (cCE), and the slope of the Constant Error (sCE) as dependent variable. Abbreviations: S = Serial Order of *t_p_* (1..10); K = Motion Kinematics (Acceleration | Deceleration); D = Motion Direction (upward | downward | leftward | rightward); P = Flashing metronome position (up | down | left | right); S-K = Static-Kinetic conditions (Acceleration | Deceleration | Flashing); G = Gravity conditions (natural g | non-natural g). In C, the Static-Kinetic model includes at the bottom the results of paired t-tests between Flashing (F), Deceleration(D) and Acceleration (A) conditions.

A. Moving Metronomes
| Factor (df) | cTV |  | sTV |  | cCE |  | sCE |  |
| --- | --- | --- | --- | --- | --- | --- | --- | --- |
|  | f-value | p-value | f-value | p-value | f-value | p-value | f-value | p-value |
| S (9) | **10.37** | **< 0.001** | 2.38 | 0.06 | **3.54** | **0.04** | **23.37** | **< 0.001** |
| K (1) | **48.65** | **< 0.001** | 0.12 | 0.73 | **9.79** | **0.01** | **13.60** | **0.00** |
| D (3) | 0.42 | 0.69 | 0.36 | 0.66 | 0.94 | 0.41 | 1.62 | 0.22 |
| SxK (9) | **5.01** | **0.00** | 0.98 | 0.44 | 2.10 | 0.10 | 2.05 | 0.09 |
| SxD (27) | 0.81 | 0.60 | 0.76 | 0.63 | 1.01 | 0.43 | 1.23 | 0.30 |
| KxD (3) | 2.22 | 0.12 | 0.43 | 0.66 | 1.45 | 0.25 | **9.61** | **0.00** |
| SxKxD (27) | 1.11 | 0.36 | 0.78 | 0.60 | 1.52 | 0.16 | 1.27 | 0.26 |

B. Flashing Metronomes
| Factor (df) | cTV |  | sTV |  | cCE |  | sCE |  |
| --- | --- | --- | --- | --- | --- | --- | --- | --- |
|  | f-value | p-value | f-value | p-value | f-value | p-value | f-value | p-value |
| S (9) | **13.35** | **< 0.001** | **2.80** | **0.03** | **10.36** | **< 0.001** | **21.75** | **< 0.001** |
| D (3) | 1.22 | 0.31 | 2.87 | 0.06 | 0.14 | 0.90 | 0.25 | 0.78 |
| SxD (27) | 1.34 | 0.23 | 1.19 | 0.31 | 1.67 | 0.13 | 0.86 | 0.54 |

C. Static-Kinetic model
| Factor (df) | cTV | sTV | cCE | sCE |
| --- | --- | --- | --- | --- |
|  | S-K (2) | S-K (2) | S-K (2) | S-K (2) |
| f-value | **46.09** | 0.78 | **44.16** | **13.53** |
| p-value | **< 0.001** | 0.46 | **< 0.001** | **< 0.001** |
| F vs D | **< 0.001** | 1.00 | **< 0.001** | **< 0.001** |
| F vs A | 0.75 | 0.61 | **< 0.001** | 1.00 |
| D vs A | **< 0.001** | 1.00 | **0.02** | **0.01** |

| Factor (df) | cTV |  | sTV |  | cCE |  | sCE |  |
| --- | --- | --- | --- | --- | --- | --- | --- | --- |
|  | f-value | p-value | f-value | p-value | f-value | p-value | f-value | p-value |
| S (9) | **7.69** | **< 0.001** | **2.23** | 0.08 | **3.22** | **0.05** | **18.78** | **< 0.001** |
| G (1) | 3.28 | 0.09 | 0.19 | 0.67 | **16.83** | **< 0.001** | 1.15 | 0.30 |
| SxG (9) | 0.65 | 0.68 | 0.38 | 0.88 | 1.71 | 0.15 | 0.94 | 0.46 |

**Table S2.** Results of the four repeated measures ANOVA models (A-D) applied to mean tap asynchronies and asynchrony variabilities. Abbreviations: S = Serial Order of taps in the Synchronization phase (1..6); Td = Target duration (450 | 550 | 650 | 750 | 850); K = Motion Kinematics (Acceleration | Deceleration); D = Motion Direction (upward | downward | leftward | rightward); P = Flashing targets position (up | down | left | right); S-K = Static-Kinetic conditions (Acceleration | Deceleration | Flashing); G = Gravity conditions (natural g | non-natural g). In C, the Static-Kinetic model includes at the bottom the results of paired t-tests between Flashing (F), Deceleration(D) and Acceleration (A) conditions.

A. Moving Metronomes
| Factor (df) | mean asynchronies |  | SD asynchronies |  |
| --- | --- | --- | --- | --- |
|  | f-value | p-value | f-value | p-value |
| S (5) | 3.45 | 0.057 | **26.75** | **< 0.001** |
| K (1) | **135.1** | **< 0.001** | **48.72** | **< 0.001** |
| D (3) | 1.39 | 0.26 | 0.47 | 0.68 |
| T<sub>d</sub> (4) | **34.95** | **< 0.001** | 2.23 | 0.11 |
| SxT<sub>d</sub> (20) | **23.22** | **< 0.001** | **4.43** | **0.002** |
| SxK (5) | **15.49** | **< 0.001** | 3.5 | 0.056 |
| SxD (15) | 1.04 | 0.38 | 0.31 | 0.89 |
| KxD (3) | 0.37 | 0.71 | 0.47 | 0.65 |
| KxT<sub>d</sub> (4) | **45.42** | **< 0.001** | 0.91 | 0.41 |
| DxT<sub>d</sub> (12) | 0.96 | 0.96 | 1.21 | 0.31 |

B. Flashing Metronomes
| Factor (df) | mean asynchronies |  | SD asynchronies |  |
| --- | --- | --- | --- | --- |
|  | f-value | p-value | f-value | p-value |
| S (5) | **57.63** | **< 0.001** | **6.45** | **0.006** |
| P (3) | 0.38 | 0.68 | 0.36 | 0.67 |
| T<sub>d</sub> (4) | **40.35** | **< 0.001** | **6.38** | **0.003** |
| SxT<sub>d</sub> (44) | 3.11 | 0.053 | 1.23 | 0.29 |
| SxP (15) | 0.96 | 0.43 | 1.18 | 0.32 |
| PxT<sub>d</sub> (12) | 0.9 | 0.47 | 0.65 | 0.66 |

C. Static-Kinetic model
| Factor (df) | mean asynchronies | SD asynchronies |
| --- | --- | --- |
|  | S-K (2) | S-K (2) |
| f-value | **82.08** | **32.44** |
| p-value | **<0.001** | **<0.001** |
| F vs D | **<0.001** | **<0.001** |
| F vs A | 0.38 | **0.005** |
| D vs A | **<0.001** | **<0.001** |

| Factor (df) | mean asynchronies |  | SD asynchronies |  |
| --- | --- | --- | --- | --- |
|  | f-value | p-value | f-value | p-value |
| S (5) | **3.89** | **0.04** | **21.75** | **< 0.001** |
| G (1) | 1.88 | 0.19 | 0.39 | 0.54 |
| T<sub>d</sub> (4) | **25.63** | **< 0.001** | 0.5 | 0.56 |
| SxG (5) | 1.91 | 0.15 | **3.61** | **0.04** |
| T<sub>d</sub>xG (4) | 1.62 | 0.2 | 0.41 | 0.68 |
| SxT<sub>d</sub> (20) | **16.44** | **< 0.001** | **2.58** | **0.02** |

**Table S3.** Results of four repeated measures ANOVA models (A-D) applied to the five Bayesian model coefficients (σm, wp, μt, σt and λ) as dependent variables. Abbreviations: S = Serial Order of *t_p_* (1,…,10); K = Motion Kinematics (Acceleration | Deceleration); D = Motion Direction (upward | downward | leftward | rightward); P = Flashing targets position (up | down | left | right); S-K = Static-Kinetic conditions (Acceleration | Deceleration | Flashing); G = Gravity conditions (natural g | non natural g). In C, the Static-Kinetic model includes at the bottom the results of paired t-tests between Flashing (F), Deceleration(D) and Acceleration (A) conditions.

A. Moving Metronomes
| Factor (df) | $\sigma_m$ | | $w_p$ | | $\mu_t$ | | $\sigma_t$ | | $\lambda$ | |
| --- | --- | --- | --- | --- | --- | --- | --- | --- | --- | --- |
|  | f-value | p-value | f-value | p-value | f-value | p-value | f-value | p-value | f-value | p-value |
| S (9) | **17.30** | **< 0.001** | **4.04** | **0.00** | **2.49** | **0.05** | **11.48** | **< 0.001** | **98.33** | **< 0.001** |
| K (1) | **34.34** | **< 0.001** | **4.73** | **0.05** | **6.17** | **0.03** | **0.66** | 0.43 | **15.78** | **0.00** |
| D (3) | 0.91 | 0.42 | 0.39 | 0.73 | 1.92 | 0.15 | 1.19 | 0.32 | 0.83 | 0.46 |
| SxK (9) | **3.44** | **0.01** | 1.78 | 0.14 | 1.80 | 0.13 | 0.61 | 0.70 | 1.56 | 0.18 |
| SxD (27) | 1.16 | 0.33 | 1.01 | 0.43 | 1.24 | 0.28 | 1.10 | 0.36 | 0.70 | 0.70 |
| KxD (3) | 0.20 | 0.84 | 1.36 | 0.27 | 1.25 | **0.30** | **5.61** | **0.01** | 1.63 | 0.20 |
| SxKxD (27) | 0.89 | 0.53 | 1.05 | 0.41 | 0.99 | 0.45 | 1.18 | 0.31 | 1.16 | 0.32 |

B. Flashing Metronomes
| Factor (df) | $\sigma_m$ | | $w_p$ | | $\mu_t$ | | $\sigma_t$ | | $\lambda$ | |
| --- | --- | --- | --- | --- | --- | --- | --- | --- | --- | --- |
|  | f-value | p-value | f-value | p-value | f-value | p-value | f-value | p-value | f-value | p-value |
| S (9) | **8.81** | **< 0.001** | **4.67** | **0.00** | **5.43** | **< 0.001** | **7.86** | **< 0.001** | **36.90** | **< 0.001** |
| D (3) | **3.73** | **0.04** | 2.50 | 0.10 | 0.66 | 0.55 | 1.20 | 0.32 | 0.66 | 0.57 |
| SxD (27) | 1.62 | 0.13 | 1.15 | 0.33 | 0.92 | 0.51 | 0.88 | 0.53 | 0.93 | 0.50 |

C. Static-Kinetic model
| Factor (df) | $\sigma_m$ | $w_p$ | $\mu_t$ | $\sigma_t$ | $\lambda$ |
| --- | --- | --- | --- | --- | --- |
|  | S-K (2) | S-K (2) | S-K (2) | S-K (2) | S-K (2) |
| f-value | **42.12** | 3.39 | **29.52** | 0.84 | **7.16** |
| p-value | **< 0.001** | 0.07 | **< 0.001** | 0.43 | **0.00** |
| F vs D | **< 0.001** | 1.00 | **< 0.001** | 0.56 | 0.15 |
| F vs A | 0.25 | **0.01** | **< 0.001** | 1.00 | 0.49 |
| D vs A | **< 0.001** | 0.14 | 0.08 | 1.00 | **0.00** |

| Factor (df) | $\sigma_m$ | | $w_p$ | | $\mu_t$ | | $\sigma_t$ | | $\lambda$ | |
| --- | --- | --- | --- | --- | --- | --- | --- | --- | --- | --- |
|  | f-value | p-value | f-value | p-value | f-value | p-value | f-value | p-value | f-value | p-value |
| S (9) | **7.75** | **< 0.001** | **2.74** | **0.03** | 1.90 | 0.12 | **7.88** | **< 0.001** | **54.37** | **< 0.001** |
| G (1) | 0.70 | 0.42 | 1.50 | 0.24 | **6.17** | **0.03** | 0.10 | 0.76 | 2.50 | 0.13 |
| SxG (9) | 0.61 | 0.66 | 0.48 | 0.79 | 0.29 | 0.93 | 0.15 | 0.98 | 0.68 | 0.65 |

## Notes

### Competing Interest Statement

The authors have declared no competing interest.

## References

1. Balasubramaniam R, Haegens S, Jazayeri M, Merchant H, Sternad D, Song JH (2021) Neural Encoding and Representation of Time for Sensorimotor Control and Learning. Journal of Neuroscience 41:866–872 Available at: https://www.jneurosci.org/content/41/5/866 [Accessed January 17, 2023].

2. Bartolo R, Prado L, Merchant H (2014) Information Processing in the Primate Basal Ganglia during Sensory-Guided and Internally Driven Rhythmic Tapping. The Journal of Neuroscience 34:3910–3923.

3. Betancourt A, Pérez O, Gámez J, Mendoza G, Merchant H (2022) Premotor population dynamics as neural substrate for auditory and visual rhythmic entrainment. bioRxiv.

4. Bingham G, Bishop R, Brody M, Bromley D, Clark E (Toby), Cooper W, Costanza R, Hale T, Hayden G, Kellert S, Norgaard R, Norton B, Payne J, Russell C, Suter G (1995) Issues in ecosystem valuation: improving information for decision making. Ecological Economics 14:73–90.

5. Burger B, Thompson MR, Luck G, Saarikallio SH, Toiviainen P (2014) Hunting for the beat in the body: On period and phase locking in music-induced movement. Front Hum Neurosci 8:903.

6. Cavada C, Goldman-Rakic PS (1993) Chapter 12 Multiple visual areas in the posterior parietal cortex of primates. Prog Brain Res 95:123–137.

7. Ceccarelli F, La Scaleia B, Russo M, Cesqui B, Gravano S, Mezzetti M, Moscatelli A, d’Avella A, Lacquaniti F, Zago M (2018) Rolling motion along an incline: Visual sensitivity to the relation between acceleration and slope. Front Neurosci 12:406.

8. Chen Y, Repp BH, Patel AD (2002) Spectral decomposition of variability in synchronization and continuation tapping: Comparisons between auditory and visual pacing and feedback conditions. Hum Mov Sci 21:515–532.

9. Cicchini GM, Arrighi R, Cecchetti L, Giusti M, Burr DC (2012) Optimal encoding of interval timing in expert percussionists. Journal of Neuroscience 32:1056–1060.

10. Collier GL, Ogden RT (2004) Adding drift to the decomposition of simple isochronous tapping: An extension of the Wing-Kristofferson model. J Exp Psychol Hum Percept Perform 30.

11. Comstock DC, Hove MJ, Balasubramaniam R (2018) Sensorimotor synchronization with auditory and visual modalities: Behavioral and neural differences. Front Comput Neurosci 12:53.

12. Crowe DA, Zarco W, Bartolo R, Merchant H (2014) Dynamic Representation of the Temporal and Sequential Structure of Rhythmic Movements in the Primate Medial Premotor Cortex. The Journal of Neuroscience 34:11972–11983.

13. Deliège I, Davidson J, Sloboda JA (2011) Music and the mind: Essays in honour of John Sloboda. Oxford University Press.

14. Delle Monache S, Indovina I, Zago M, Daprati E, Lacquaniti F, Bosco G (2021) Watching the Effects of Gravity. Vestibular Cortex and the Neural Representation of “Visual” Gravity. Front Integr Neurosci 15.

15. Delle Monache S, Lacquaniti F, Bosco G (2017) Differential contributions to the interception of occluded ballistic trajectories by the temporoparietal junction, area hMT/V5+, and the intraparietal cortex. J Neurophysiol 118:1809–1823.

16. Delle Monache S, Lacquaniti F, Bosco G (2019) Ocular tracking of occluded ballistic trajectories: Effects of visual context and of target law of motion. J Vis 19:13–13.

17. Fraisse P (1963) The psychology of time. Harper & Row.

18. Fraisse P (1978) TIME AND RHYTHM PERCEPTION. Perceptual Coding:203–254.

19. Gámez J, Mendoza G, Prado L, Betancourt A, Merchant H (2018) The amplitude in periodic neural state trajectories underlies the tempo of rhythmic tapping. Enviado.

20. Gámez J, Mendoza G, Prado L, Betancourt A, Merchant H (2019) The amplitude in periodic neural state trajectories underlies the tempo of rhythmic tapping. PLoS Biol 17:e3000054.

21. Gan L, Huang Y, Zhou L, Qian C, Wu X (2015) Synchronization to a bouncing ball with a realistic motion trajectory. Scientific Reports 2015 5:1 5:1–9.

22. García-Garibay O, Cadena-Valencia J, Merchant H, de Lafuente V (2016) Monkeys share the human ability to internally maintain a temporal rhythm. Front Psychol 7:1971.

23. García-Saldivar P, de León C, Concha L, Merchant H (2022) White matter structural bases for predictive tapping synchronization. bioRxiv.

24. Gibbon J, Malapani C, Dale CL, Gallistel CR (1997) Toward a neurobiology of temporal cognition: advances and challenges. Curr Opin Neurobiol 7:170–184.

25. Globerson E, Flash T, Eitan Z, Globerson E, Flash T, Eitan Z (2021) Space, Time and Expression in Orchestral Conducting. :199–212 Available at: https://link.springer.com/chapter/10.1007/978-3-030-57227-3_10 [Accessed January 17, 2023].

26. Grahn JA, Rowe JB (2009) Feeling the Beat: Premotor and Striatal Interactions in Musicians and Nonmusicians during Beat Perception. Journal of Neuroscience 29:7540–7548.

27. Guttman SE, Gilroy LA, Blake R (2005) Hearing what the eyes see: Auditory encoding of visual temporal sequences. Psychol Sci 16:228–235.

28. Honing H (2012) Without it no music: beat induction as a fundamental musical trait. Ann N Y Acad Sci 1252:85–91.

29. Honing H, Merchant H (2014) Differences in auditory timing between human and nonhuman primates. Behavioral and Brain Sciences 37:557–558 Available at: https://www.cambridge.org/core/journals/behavioral-and-brain-sciences/article/abs/differences-in-auditory-timing-between-human-and-nonhuman-primates/B32ED923A8E60E2F29AC566DD29DFDBD [Accessed February 3, 2023].

30. Honing H, Merchant H, Háden GP, Prado L, Bartolo R (2012) Rhesus Monkeys (Macaca mulatta) Detect Rhythmic Groups in Music, but Not the Beat. PLoS One 7:e51369.

31. Hove MJ, Iversen JR, Zhang A, Repp BH (2013) Synchronization with competing visual and auditory rhythms: Bouncing ball meets metronome. Psychol Res 77:388–398.

32. Hove MJ, Spivey MJ, Krumhansl CL (2010) Compatibility of Motion Facilitates Visuomotor Synchronization. J Exp Psychol Hum Percept Perform 36:1525–1534.

33. Indovina I, Maffei V, Bosco G, Zago M, Macaluso E, Lacquaniti F (2005) Representation of Visual Gravitational Motion in the Human Vestibular Cortex. Science (1979) 308:416–419.

34. Iversen JR, Patel AD, Nicodemus B, Emmorey K (2015) Synchronization to auditory and visual rhythms in hearing and deaf individuals. Cognition 134:232–244.

35. Ivry RB, Hazeltine RE (1995) Perception and Production of Temporal Intervals Across a Range of Durations: Evidence for a Common Timing Mechanism. J Exp Psychol Hum Percept Perform 21:3–18 Available at: /doiLanding?doi=10.1037%2F0096-1523.21.1.3 [Accessed December 8, 2022].

36. Jantzen KJ, Ratcliff BR, Jantzen MG (2017) Cortical Networks for Correcting Errors in Sensorimotor Synchronization Depend on the Direction of Asynchrony. https://doi.org/101080/0022289520171327414 50:235–248.

37. Jazayeri M, Shadlen MN (2010) Temporal context calibrates interval timing. Nat Neurosci 13:1020–1026.

38. Kim IK, Spelke ES (1992) Infants’ Sensitivity to Effects of Gravity on Visible Object Motion. J Exp Psychol Hum Percept Perform 18.

39. Kubovy M (1988) Should We Resist the Seductiveness of the Space: Time:: Vision:Audition Analogy? J Exp Psychol Hum Percept Perform 14:318–320.

40. La Scaleia B, Zago M, Lacquaniti F (2015) Hand interception of occluded motion in humans: A test of model-based vs.on-line control. J Neurophysiol 114:1577–1592.

41. Lacquaniti F, Bosco G, Gravano S, Indovina I, La Scaleia B, Maffei V, Zago M (2015) Gravity in the Brain as a Reference for Space and Time Perception. Multisens Res 28:397–426.

42. Lacquaniti F, Bosco G, Indovina I, La Scaleia B, Maffei V, Moscatelli A, Zago M (2013) Visual gravitational motion and the vestibular system in humans. Front Integr Neurosci 7:101.

43. Lacquaniti F, Maioli C (1989) The role of preparation in tuning anticipatory and reflex responses during catching. Journal of Neuroscience 9:134–148.

44. Lee DN, Reddish PE (1981) Plummeting gannets: a paradigm of ecological optics. Nature 1981 293:5830 293:293–294.

45. Lenc T, Merchant H, Keller PE, Honing H, Varlet M, Nozaradan S (2021) Mapping between sound, brain and behaviour: Four-level framework for understanding rhythm processing in humans and non-human primates. Philosophical Transactions of the Royal Society B: Biological Sciences 376.

46. Li Y, Wang Y, Cui H (2022) Posterior parietal cortex predicts upcoming movement in dynamic sensorimotor control. Proc Natl Acad Sci U S A 119:e2118903119 Available at: https://www.pnas.org/doi/abs/10.1073/pnas.2118903119 [Accessed December 8, 2022].

47. Lisberger SG, Movshon JA (1999) Visual Motion Analysis for Pursuit Eye Movements in Area MT of Macaque Monkeys. Journal of Neuroscience 19:2224–2246.

48. Luck G, Nte S (2008) An investigation of conductors’ temporal gestures and conductor— musician synchronization, and a first experiment. Psychol Music 36:81–99.

49. Luck G, Sloboda JA (2009) Spatio-temporal cues for visually mediated synchronization. Music Percept 26:465–473.

50. Madison G (2001) Variability in isochronous tapping: Higher order dependencies as a function of intertap interval. J Exp Psychol Hum Percept Perform 27:411–422.

51. McAuley JD, Jones MR (2003) Modeling effects of rhythmic context on perceived duration: a comparison of interval and entrainment approaches to short-interval timing. J Exp Psychol Hum Percept Perform 29:1102–1125.

52. McAuley JD, Jones MR, Holub S, Johnston HM, Miller NS (2006) The time of our lives: Life span development of timing and event tracking. J Exp Psychol Gen 135:348–367.

53. Mendoza G, Merchant H (2014) Motor system evolution and the emergence of high cognitive functions. Prog Neurobiol 122:73–93.

54. Merchant H, Averbeck BB (2017) The Computational and Neural Basis of Rhythmic Timing in Medial Premotor Cortex. Journal of Neuroscience 37:4552–4564 Available at: https://www.jneurosci.org/content/37/17/4552 [Accessed December 19, 2022].

55. Merchant H, Bartolo R (2018) Primate beta oscillations and rhythmic behaviors. J Neural Transm 125:461–470.

56. Merchant H, Battaglia-Mayer A, Georgopoulos AP (2003) Functional organization of parietal neuronal responses to optic-flow stimuli. J Neurophysiol 90:675–682 Available at: https://journals.physiology.org/doi/10.1152/jn.00331.2003 [Accessed December 8, 2022].

57. Merchant H, Battaglia-Mayer A, Georgopoulos AP (2004a) Neural Responses during Interception of Real and Apparent Circularly Moving Stimuli in Motor Cortex and Area 7a. Cerebral Cortex 14:314–331 Available at: https://academic.oup.com/cercor/article/14/3/314/418454 [Accessed December 8, 2022].

58. Merchant H, Battaglia-Mayer A, Georgopoulos AP (2004b) Neural responses in motor cortex and area 7a to real and apparent motion. Exp Brain Res 154:291–307 Available at: https://link.springer.com/article/10.1007/s00221-003-1664-5 [Accessed February 3, 2023].

59. Merchant H, Georgopoulos AP (2006) Neurophysiology of perceptual and motor aspects of interception. J Neurophysiol 95:1–13 Available at: https://journals.physiology.org/doi/10.1152/jn.00422.2005 [Accessed December 8, 2022].

60. Merchant H, Grahn J, Trainor L, Rohrmeier M, Fitch WT (2015a) Finding the beat: a neural perspective across humans and non-human primates. Philosophical Transactions of the Royal Society B: Biological Sciences 370.

61. Merchant H, Honing H (2014) Are non-human primates capable of rhythmic entrainment? Evidence for the gradual audiomotor evolution hypothesis. Front Neurosci 7:274.

62. Merchant H, Luciana M, Hooper C, Majestic S, Tuite P (2008a) Interval timing and Parkinson’s disease: heterogeneity in temporal performance. Exp Brain Res 184:233– 248 Available at: http://dx.doi.org/10.1007/s00221-007-1097-7.

63. Merchant H, Perez O, Bartolo R, Méndez JC, Mendoza G, Gámez J, Yc K, Prado L (2015b) Sensorimotor neural dynamics during isochronous tapping in the medial premotor cortex of the macaque. European Journal of Neuroscience 41:586–602.

64. Merchant H, Perez O, Zarco W, Gámez J (2013) Interval Tuning in the Primate Medial Premotor Cortex as a General Timing Mechanism. The Journal of Neuroscience 33:9082–9096.

65. Merchant H, Zarco W, Perez O, Prado L, Bartolo R (2011) Measuring time with different neural chronometers during a synchronization-continuation task. Proceedings of the National Academy of Sciences 108:19784–19789 Available at: http://www.pnas.org/content/108/49/19784.abstract.

66. Merchant H, Zarco W, Prado L (2008b) Do We Have a Common Mechanism for Measuring Time in the Hundreds of Millisecond Range? Evidence From Multiple-Interval Timing Tasks. J Neurophysiol 99:939–949.

67. Moscatelli A, Lacquaniti F (2011) The weight of time: Gravitational force enhances discrimination of visual motion duration. J Vis 11:5–5.

68. Movshon JA, Lisberger SG, Krauzlis RJ (1990) Visual Cortical Signals Supporting Smooth Pursuit Eye Movements. Cold Spring Harb Symp Quant Biol 55:707–716.

69. Patel AD, Iversen JR (2014) The evolutionary neuroscience of musical beat perception: The Action Simulation for Auditory Prediction (ASAP) hypothesis. Front Syst Neurosci 8:57.

70. Perez O, Merchant H (2018) The Synaptic Properties of Cells Define the Hallmarks of Interval Timing in a Recurrent Neural Network. J Neurosci 38:4186–4199.

71. Petzschner FH, Glasauer S (2011) Iterative Bayesian Estimation as an Explanation for Range and Regression Effects: A Study on Human Path Integration. Journal of Neuroscience 31:17220–17229.

72. Petzschner FH, Glasauer S, Stephan KE (2015) A Bayesian perspective on magnitude estimation. Trends Cogn Sci 19:285–293.

73. Pittenger JB (1990) Detection of Violations of the Law of Pendulum Motion: Observers’ Sensitivity to the Relation Between Period and Length. Ecological Psychology 2.

74. Price NSC, Ono S, Mustari MJ, Ibootson MR (2005) Comparing acceleration and speed tuning in macaque MT: Physiology and modeling. J Neurophysiol 94:3451–3464.

75. Rajendran VG, Teki S, Schnupp JWH (2018) Temporal Processing in Audition: Insights from Music. Neuroscience 389:4–18.

76. Repp BH (2005) Sensorimotor synchronization: A review of the tapping literature. Psychonomic Bulletin & Review 2005 12:6 12:969–992.

77. Repp BH, Penel A (2002) Auditory Dominance in Temporal Processing: New Evidence from Synchronization with Simultaneous Visual and Auditory Sequences. J Exp Psychol Hum Percept Perform 28:1085–1099.

78. Repp BH, Penel A (2004) Rhythmic movement is attracted more strongly to auditory than to visual rhythms. Psychol Res 68:252–270.

79. Sánchez-Moncada I, Concha L, Merchant H (2022) Pre-supplementary motor cortex mediates learning transfer from perceptual to motor timing. bioRxiv.

80. Schlack A, Albright TD (2007) Remembering Visual Motion: Neural Correlates of Associative Plasticity and Motion Recall in Cortical Area MT. Neuron 53:881–890.

81. Schramm R, Jung CR, Miranda ER (2015) Dynamic time warping for music conducting gestures evaluation. IEEE Trans Multimedia 17:243–255.

82. Toh LW, Chao W, Chen YS (2013) An interactive conducting system using Kinect. Proc (IEEE Int Conf Multimed Expo).

83. Twardy CR, Bingham GP (2002) Causation, causal perception, and conservation laws. Perception & Psychophysics 2002 64:6 64:956–968.

84. Welch RB, Warren DH (1980) Immediate perceptual response to intersensory discrepancy. Psychol Bull 88.

85. Wing AM (2002) Voluntary Timing and Brain Function: An Information Processing Approach. Brain Cogn 48:7–30.

86. Wing AM, Kristofferson AB (1973a) The timing of interresponse intervals. Perception & Psychophysics 1973 13:3 13:455–460.

87. Wing AM, Kristofferson AB (1973b) Response delays and the timing of discrete motor responses. Perception & Psychophysics 1973 14:1 14:5–12.

88. Woodrow H (1934) The temporal indifference interval determined by the method of mean error. J Exp Psychol 17:167–188.

89. Zarco W, Merchant H, Prado L, Mendez JC (2009) Subsecond Timing in Primates: Comparison of Interval Production Between Human Subjects and Rhesus Monkeys. J Neurophysiol 102:3191–3202 Available at: http://jn.physiology.org/content/102/6/3191.abstract.

